# Computationally accelerated identification of P-glycoprotein inhibitors

**DOI:** 10.1101/2024.03.05.583428

**Authors:** Lauren A. McCormick, James W. McCormick, Chanyang Park, Courtney A. Follit, John G. Wise, Pia D. Vogel

**Affiliations:** Department of Biological Sciences, Southern Methodist University, Dallas, Texas, United States; Center for Drug Discovery, Design and Delivery, Southern Methodist University, Dallas, Texas, United States; Center for Scientific Computation, Southern Methodist University, Dallas, Texas

## Abstract

Overexpression of the polyspecific efflux transporter, P-glycoprotein (P-gp, MDR1, *ABCB1*), is a major mechanism by which cancer cells acquire multidrug resistance (MDR), the resistance to diverse chemotherapeutic drugs. Inhibiting drug transport by P-gp can resensitize cancer cells to chemotherapy, but there are no P-gp inhibitors available to patients. Clinically unsuccessful P-gp inhibitors tend to bind at the pump’s transmembrane drug binding domains and are often P-gp transport substrates, resulting in lowered intracellular concentration of the drug and altered pharmacokinetics. In prior work, we used computationally accelerated drug discovery to identify novel P-gp inhibitors that target the pump’s cytoplasmic nucleotide binding domains. Our first-draft study provided conclusive evidence that the nucleotide binding domains of P-gp are viable targets for drug discovery. Here we develop an enhanced, computationally accelerated drug discovery pipeline that expands upon our prior work by iteratively screening compounds against multiple conformations of P-gp with molecular docking. Targeted molecular dynamics simulations with our homology model of human P-gp were used to generate docking receptors in conformations mimicking a putative drug transport cycle. We offset the increased computational complexity using custom Tanimoto chemical datasets, which maximize the chemical diversity of ligands screened by docking. Using our expanded, virtual-assisted pipeline, we identified nine novel P-gp inhibitors that reverse MDR in two types of P-gp overexpressing human cancer cell lines, reflecting a 13.4% hit rate. Of these inhibitors, all were non-toxic to non-cancerous human cells, and six were not likely to be transport substrates of P-gp. Our novel P-gp inhibitors are chemically diverse and are good candidates for lead optimization. Our results demonstrate that the nucleotide binding domains of P-gp are an underappreciated target in the effort to reverse P-gp-mediated multidrug resistance in cancer.

## Introduction

Multidrug resistance (MDR) describes the ability of cells to become resistant to structurally and chemically diverse drugs [1]. Cancer cells can acquire MDR over the course of chemotherapeutic treatment, and MDR therefore is a significant obstacle to the successful treatment of human cancers [2]. Cancer cells often acquire MDR by overexpressing ATP-binding cassette (ABC) transporters [3]. ABC transporters are dynamic, ATP-powered efflux pumps that confer MDR by pumping chemotherapeutic drugs out of the respective cancer cell. This transport activity lowers the intracellular concentration of the chemotherapeutic, thereby substantially reducing the drug’s efficacy [3]. Among the most clinically relevant ABC transporters is P-glycoprotein (P-gp, MDR1, *ABCB1*), a remarkably promiscuous pump that is commonly overexpressed by multidrug resistant cancers [3, 4]. P-gp harnesses the power of ATP hydrolysis to pump chemically and structurally diverse substrates out of the cell. The pump’s cytoplasmic nucleotide binding domains (NBDs) bind and hydrolyze ATP, and its membrane-embedded drug binding domains (DBDs) recognize and extrude substrates [5]. P-gp’s substrate profile is incredibly diverse, ranging from small drugs to bulky amyloid β peptides [6], and importantly, includes a staggeringly diverse array of chemotherapeutics. Inhibiting P-gp’s efflux activity can resensitize MDR cancers to chemotherapy [7–10], but the search for clinically successful P-gp inhibitors has been fraught with failure.

Decades of work have yielded three generations of structurally and chemically diverse P-gp inhibitors [1, 11] (S1 Table). First and second generation inhibitors were often identified by chance, and as such targeted P-gp with relatively low specificity [1, 11], requiring high therapeutic doses and causing them to fail clinical trials because of off-target toxicity. All first and second-generation P-gp inhibitors are thought to bind at the DBDs, and many were also shown to be P-gp transport substrates [11, 12]. Third-generation inhibitors (tariquidar, zosuquidar, and their derivatives) improved on earlier generations by targeting the DBDs with a higher affinity, but these inhibitors still failed in clinical trials due to toxicity and off-target effects [13, 14]. It was initially thought that tariquidar and zosuquidar were not P-gp transport substrates, but later work has shown otherwise [15, 16]. In summary, while P-gp remains a clinically relevant target, efforts to discover or design a clinically effective inhibitor have been unsuccessful.

To date, most known P-gp inhibitors, even those with relatively high binding affinities like tariquidar, have been shown to bind at the DBDs. We hypothesize that this shared characteristic may contribute to the failure of these inhibitors in clinical trials. Treatment with P-gp transport substrates inherently requires a higher therapeutic dose – the pump continuously ejects its own inhibitor from the cell, and inhibition is only effective while the molecule resides within the DBDs. Thus, if a P-gp inhibitor is also a transport substrate, P-gp transport activity inherently lowers the drug’s inhibitory effect. This in turn raises the required dose of inhibitor for patient treatment and likely contributes to the adverse side effects observed in clinical trials [16].

A growing body of evidence suggests that the DBDs are no longer suitable targets for the design of small molecule inhibitors [11]. P-gp’s substrate profile is so diverse that it is challenging to find a DBD inhibitor that is not a transport substrate [6, 15, 17, 18]. The DBDs of P-gp are key to the pump’s promiscuity. The DBDs large, hydrophobic, and flexible. They are large enough to accommodate substrates like amyloid-β 42 (4500 Daltons) and to provide multiple residue contact pathways for substrates during transport [6, 19]. Combined with the failure of previous generations of P-gp inhibitors, these observations strongly indicate that the DBDs are no longer promising targets. However, P-gp’s nucleotide binding domains (NBDs) have not been thoroughly explored.

In previous studies, our group identified novel P-gp inhibitors using a computationally-assisted drug discovery program targeting the NBDs. Our early screens successfully identified three inhibitors that reverse MDR in human cancer cells with a 7% hit rate [7, 8]. A fourth compound was identified that interacts with the NBDs in assays with purified protein, but is likely excluded from cells due to its negative charge at neutral pH (6). This first-draft attempt suggested that the NBDs are suitable candidates for more intensive drug screening.

Here we built upon our earlier work by significantly expanding our pilot drug discovery pipeline. First, molecular docking was used to virtually screen millions of druglike molecules against P-gp. Our first study used only one conformation of P-gp for docking [7]. While that study was ultimately successful, we wanted to screen against a more complete sample of P-gp conformations during transport. Since high-resolution structures of human P-gp were not available when the docking presented here was performed, targeted molecular dynamics (TMD) simulations were used to generate dynamic P-gp structures for molecular docking. Millions of molecules were then iteratively screened against the NBDs and DBDs of each target, with the goal of selecting molecules that were predicted to prefer the NBDs. A subset of molecules was tested *in vitro* for the ability to reverse MDR in two types of P-gp-overexpressing human cancer cell lines. The top hits were screened for inherent toxicity with a non-cancerous human cell line, assessed as P-gp transport substrates using LC-MS/MS, and tested for ability to increase retention of the P-gp substrate and chemotherapeutic daunorubicin.

Using a combination of targeted MD simulations, molecular docking, and cell-based assays, we identified nine novel P-gp inhibitors that reverse MDR, reflecting a 13.4% hit rate. These inhibitors were effective against two types of human P-gp overexpressing cancer lines: DU145-TXR, a prostate cancer line, and A2780-ADR, an ovarian cancer line. These nine inhibitors were found to be non-toxic to non-cancerous human cells, and six were then shown to be unlikely transport substrates of P-gp. Four of this final molecule set were found to enhance the intracellular accumulation of daunorubicin. In summary, these nine novel P-gp inhibitors are good candidates for lead optimization and further testing. This work reinforces the idea that the NBDs are promising targets in the fight against multidrug resistance in cancer. This ‘enhanced screening’ approach can be adapted for drug discovery screens that target other clinically relevant ABC transporters such as breast cancer resistance protein (BCRP, *ABCG2*) and MsbA from *Escherichia coli*.

## Materials and methods

### Receptor generation for virtual screens against human P-gp

When these docking studies were performed, the available structures of P-gp were extremely limited in conformational diversity and resolution, and the highest-resolution structures were primarily of human P-gp homologues. Molecular dynamics simulations with a homology model of human P-gp were used to overcome this limitation and generate structures of P-gp in diverse conformations as follows. Using the homology model of human P-gp from our prior work [20], a putative catalytic cycle of P-glycoprotein (P-gp, MDR1, *ABCB1*) was simulated using Targeted Molecular Dynamics (TMD) simulations with NAMD [21], using procedures as described in [19, 21].

Briefly, using VMD[22] our human P-gp model was inserted into a POPC phospholipid bilayer, solvated with TIP3P water, and neutralized with Na^+^ and Cl^-^ counterions, as in [19]. Nucleotides (ATP) and Mg^2+^ ions were added into the nucleotide binding domains of P-gp. MD simulations were performed at 310 K (37 °C) with Langevin temperature and pressure control, particle-mesh Ewald electrostatics, and constant temperature and pressure (*NPT*) in a periodic cell, as in [19]. For TMD, target structures were aligned with the P-gp model using STAMP [23] from the Multiple Alignment [24] module included in VMD [22]. The coordinates of homologous pairs of alpha carbon (Cα) atoms (paired Cα atoms between the model and respective target) were used as target coordinates for TMD. TMD simulations were performed in NAMD, and forces were applied using in-house *Tcl* scripts that aimed to gently move the atoms toward the target coordinates, as used by us previously in [6, 19]. Force magnitude is calculated to be inversely proportional to the Root Mean Squared Deviation (RMSD) of the distance between the paired Cα atoms of model and target.

The following structures of P-gp homologues were used as target structures for TMD: an open-to-the-inside mouse P-gp structure with wide separation between the NBDs (PDB entry 4KSB[25]); an open-to-the-inside structure of the *E. coli* homologue MsbA with lesser NBD separation than 4KSB (PDB entry 3B5X [26]); closed-to-the-inside structure of MsbA with NBDs engaged (PDB entry 3B5Z [26]); an closed-to-the-inside structure of the Sav1866 transporter from *Staphylococcus aureus* with NBDs engaged (PDB entry 2HYD [27]). It is important to note that these simulations, and subsequent molecular docking studies, were performed before high-resolution structures of P-gp, and indeed before any structures of human P-gp, were available for use.

TMD simulations were performed sequentially to guide our human P-gp model into the conformation of each structure, with the goal of mimicking the conformational transitions of a putative catalytic efflux cycle, and ultimately of producing human P-gp conformations for molecular docking screens. From TMD simulations, we isolated ten distinct conformations of P-gp for use in molecular docking studies: an approximate 4KSB position, 3B5X position, two 3B5Z positions, two 2HYD positions, and two positions between 4KSB and 3B5X named the “Transition” position (S1 – S2 Fig). Three structures (4KSB_DBD, Transition_DBD, and 3B5X_DBD) (S1 Fig, Boxes A – C, S2 Fig boxes A – C) were used for docking against the drug binding domains (DBD), and five (3B5Z_NBD_1, 3B5Z_NBD_2, 2HYD_NBD_1, 2HYD_NBD_2 and Transition NBD) were used for docking against the nucleotide binding domains (NBD) (S1 Fig, Boxes D – F; S2 Fig Boxes D – E). The NBD search areas shown in S1 Fig D – F were divided into two separate dock boxes, each targeting the respective individual NBD. Receptor files were prepared as PDBQTs for docking using AutoDock Tools [28].

### Ligand selection and preparation for docking

Compounds were taken from the ZINC12 Clean Drug-Like Subset of 13,195,609 molecules (31). Molinspiration (*mib)* software was used to calculate logP, polar surface area, molecular weight, number of hydrogen-bond donors and acceptors and number of rotatable bonds for each ligand [29]. To predict the aqueous solubility of our experimental compounds, the AMSOL program was used to calculate polar and apolar desolvation energies [30]. The criteria for ligand selection conformed to Lipinski-Veber rule for characteristics of drug-like or therapeutic molecules with oral bioavailability [31, 32]: (1) a molecular weight less than 500 and greater than 150 Daltons; (2) an octanol-water partition coefficient (log P) less than 5; (3) a topological polar surface area less than 150 Å^2^; (4) less than 5 hydrogen donors; and (5) less than 10 hydrogen acceptors. The number of rotatable bonds was less-than or equal-to 7, and the “clean” designator indicates that aldehydes and thiols are filtered out.

Tanimoto sets [33] are used to generate subsets of drug-like molecules that represent the ‘chemical diversity’ of the total molecule dataset. First, each molecule in the original dataset is given a unique fingerprint. The components of the fingerprint indicate the presence or absence of specific molecule fragments (such as chemical groups). Then, the Tanimoto algorithm computes the similarity between each molecule in the dataset, with the goal of identifying a subset of unique molecules that represent the chemical diversity in the dataset. The chemical diversity is expressed as a percentage (e.g. 90%), which indicates that the corresponding Tanimoto set contains 90% of the chemical diversity in that dataset. The lower the percentage, the fewer compounds in the Tanimoto set. Dataset size becomes very relevant when considering molecular docking against multiple receptors, like the docking screens presented in this study.

The dataset used to identify compounds SMU 58-68 was obtained from a pre-calculated 90% Tanimoto cutoff set containing 123,510 “Clean Drug-Like Compounds” from the Zinc^12^ database [34]. Ligands from the pre-calculated 90% Tanimoto set were protonated according to a pH of 7. We found that this subset was not structurally diverse, especially in molecular weight, and therefore only 10 compounds were purchased from it. For compounds SMU 68 – 124, we generated a custom 95% Tanimoto set from the ZINC Clean Drug-Like Subset using cactvs (http://www.xemistry.com/) and SUBSET (https://cactus.nci.nih.gov/subset/); this resulted in a final dataset of 158,000 diverse molecules, protonated at pH of 7.

Taken together, the ligands that were selected from Tanimoto sets of the ZINC database encompassed 158,000 compounds in total. In preparation for docking, all ligands (pulled as mol2 files from the ZINC database) were converted to the AutoDock-compatible PDBQT format using Open Babel [35]. Open Babel 2.3.2 was used for the Zinc12 90% Tanimoto dataset and Open Babel 2.4.0 was used for the custom 95% Tanimoto dataset. Open Babel conversion protonates ligands according to specified pH; ligands were protonated to a pH of 7.

### Ligand docking and iterative screening with *AutoDock Vina*

AutoDock Vina was used to dock each molecule against each target, totaling over 4.2 million individual docking experiments. Docking experiments AutoDock Vina and the generation of dock boxes with AutoDock 4 were performed as described in [36]. Briefly, protein receptor files were converted to PDBQT with AutoDock 4.2. Dock boxes were designed to encompass the ligand-binding regions of the DBDs, but included the full structure of the NBDs. Docking grids were created with AutoDock; the placement and size of all boxes are shown in S1 – S2 Fig and in S2 Table. Docking experiments with AutoDock Vina used default settings (9 binding modes, maximum energy difference of 3 kcal/mol) with the exception of exhaustiveness, which was set to 128 instead of the default of 8. AutoDock Vina’s scoring function is described in detail in [36]. AutoDock Vina ranks resultant ligand docking poses by predicted affinities/binding energies, and outputs *estimated* binding energy (ΔG, kcal/mol). For each ligand-target combination, we extracted the docking pose with the best estimated binding energy (ΔG, kcal/mol) and used that estimated energy to calculate an *estimated* equilibrium constant of dissociation (K_D_). Estimated K_D_ was calculated at 310 K, to match the temperature of MD simulations, using the quantitative relationship between K_D_ and Gibbs free energy (ΔG),

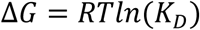

where “T” is the temperature in degrees kelvin (310 K for our work), R is the gas constant with 1.987 x 10^-3^ kcal mol^-1^ K^-1^, and ΔG is estimated free energy of binding (derived from docking calculation) in kcal mol^-1^. We emphasize that the docking-derived binding energies and affinities were treated purely as estimates, for use primarily to calculate a ratio of predicted preference for the NBDs of P-gp over the DBDs. Actual measures of binding affinities to either region are outside the scope of this work.

The goal of the docking studies was to identify molecules that were *predicted* to prefer the NBDs instead of the DBDs. To make this prediction, the ratio of *estimated* binding affinities was calculated as follows: K_D_^DBD^ / K_D_^NBD^. The smaller the value of K_D_, the tighter the affinity, and vice versa. Thus, a molecule with a large estimated K_D_^DBD^ (weak binding) and a small estimated K_D_^NBD^ (tight binding) will have a large ratio. A ligand’s respective ratio for each receptor pair (e.g., K_D_^NBD of 3B5X^ / K_D_^DBD of 3B5X^) was calculated and used to select 100 ligands as ‘top hits’ of this first round of screening.

To perform iterative screening, the 100 ligands with the best DBD/NBD ratio were used to find molecules in the original ZINC12 Clean, Drug-like database with 99-70% chemical similarity. This approach produced a second dataset of ∼100 - 10,000 molecules per top ligand. These ligand sets were subsequently docked and analyzed in the same way as the first set – by calculating the ratio of estimated affinity to DBD versus NBD. From this second iterative screen, a subset of molecules with favorable DBD/NBD ratios were selected for testing *in vitro*. Examples of ligand docking locations are shown in S3 Fig. The estimated DBD/NBD ratio of the seven molecules identified by subsequent *in vitro* screens are shown in S3 Table.

### Property prediction and selection

The top 100 ligands against each target were subjected to property prediction using the Online Chemical Database (OCHEM) [37]. Top hits converted to SMILES (simplified molecular-input line-entry system) strings and processed by OCHEM, producing predictions for logP and Solubility, Environmental toxicity, Ames test, CYP3A4 inhibition, CYP2D6 inhibition, CYP2C19 inhibition, CYP2C9 inhibition, CYP1A2 inhibition, Melting Point best (Estate), Pyrolysis OEstate submodel, Water solubility (GSE) based on logP and Melting Point, ALOGPS 2.1 logS, ALOGPS 2.1 logP, DMSO solubility, and any known toxic effects from literature. Along with the specific docking location to the receptor as visualized in VMD, the structural, chemical, and docking information was used to select ligands for purchase and subsequent testing.

### Cell lines and cell culture

The chemotherapeutic sensitive DU145 human prostate cancer cells and the multidrug resistant sub-line, DU145-TXR, were generous gifts from Dr. Evan Keller (University of Michigan, Ann Arbor, MI) [38]. The MDR, P-gp overexpressing DU145-TXR cells were maintained under positive selection pressure by supplementing complete medium with 10 nM paclitaxel (Acros Organics, NJ). In addition to the aforementioned cell lines, the chemotherapeutic sensitive A2780 ovarian cancer cells (93112519, Sigma) and the multidrug resistant A2780-ADR (93112520, Sigma) were maintained in complete RPMI media consisting of RPMI-1640 with L-glutamine, 10% fetal bovine serum (FBS; Corning), 100 U/mL penicillin and 100 μg/mL streptomycin in a humidified incubator at 37°C and 5% CO2. The MDR, P-gp overexpressing A2780-ADR cell line was maintained under positive selection pressure by supplementing complete medium with 100 nM doxorubicin (Fisher Scientific, NJ). The non-cancerous HFL1 (human lung fibroblast) cell line cell line was a generous gift from Dr. Robert Harrod (Southern Methodist University, Dallas, TX) and was maintained in complete F12K media consisting of F12K with 10% fetal bovine serum, 100 U/mL penicillin, and 100 µg/mL streptomycin (Gibco). Flasks and 96 well plates that were used with the non-cancerous line HFL1 were pre-treated with 0.1 mg/ml Collagen Type 1 (Gibco) and rinsed with PBS (Fisher).

### Testing of experimental compounds in cell culture

Culture of the DU145, DU145-TXR, and HFL1 cell lines was performed as described in [8]. Culture of the A2780 and A2780-ADR cell lines was performed as described in [9]. MTT assays were performed as described in [8] with minor modifications. LC-MS/MS accumulation assays were performed as described in [9] with some modifications. DU145-TXR cells were exposed to 500 nmol/L PTX, a dosage that has been shown to result in greater than 85% survival of the multidrug-resistant DU145-TXR cell line, and less than 10% survival of the chemotherapy-sensitive DU145 parental cell line [8]. A concentration of 15 µM was used because in prior work by us, concentrations of 25 µM were shown to robustly inhibit P-gp-catalyzed ATP hydrolysis [7]. The inhibitor concentration was lowered to 15 µM here, as the goal of the study was to identify compounds that were effective in reversing MDR at low concentrations, and to determine if the expanded virtual-assisted methods improved on our first-draft attempt. Compounds 56 through 98 were pre-screened using Resazurin cell viability assays: cells were incubated with 15 µM compound with or without 500 nM PTX for 48 hours, after which cell survival was assessed similar to [39]. Data represent the mean of two separate experiments, with n = 3 samples per experiment (n = 6 total) (S4 Table). The top performing compounds from this initial screen were re-tested with MTT viability assays against DU145-TXR cells, as described below.

### MTT cell viability assays

MTT assays measure the reduction of yellow, water soluble 3-(4,5-dimethylthiazol-2-yl)-2,5-diphenyltetrazolium bromide (MTT) to blue, insoluble formazan crystals by cellular reductase processes in living cells [40]. Cells were trypsinized from monolayers and seeded in complete medium in 96 well plates. Cancerous cell lines (DU145, DU145-TXR, A2780, A2780-ADR) were seeded at 3,000 cells per well in complete RPMI medium. The non-cancerous HFL1 line was seeded at 4,000 cells per well in complete F12K medium, in collagen-treated plates (Collagen Type 1, Gibco). Cells were incubated with experimental compounds for 48 hours at 37 °C and 5% CO_2_ in a humidified incubator. At 48 hours, MTT was added (20 µL/well of 5 mg/mL MTT in PBS) to each well. After incubation for 4 hours, the media was removed, and the formazan crystals were dissolved in 100 µL DMSO per well. Plates containing the semi-adherent A2780 or A2780-ADR lines were spun at 1400 rpm for 3 minutes prior to aspiration of media, and the addition of DMSO. Plates were shaken for 10 minutes at 500 rpm on an Orbital shaker (LabDoctor from MidSci, St Louis, MO). The absorbance was measured at 570 nm using a Bio-Tek Cytation 5 (Bio-Tek, Winooski, VT). The measured absorbance value was correlated with the number of metabolically active cells in that well, corrected for absorbance of media. The percent of cell viability (termed ‘survival’ here) was calculated as a percentage of survival of the control condition, which refers to vehicle-treated (DMSO as vehicle) cells, or vehicle-and- chemotherapeutic-treated cells, if testing a compound’s ability to re-sensitize cells to a chemotherapeutic. Percent survival was calculated using the following equation:

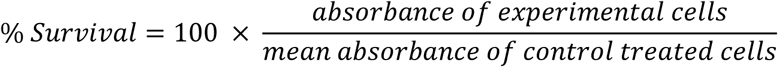

For DU145-TXR, DU145, HFL1 and A2780 cells - data are the mean ± one standard deviation, 8 samples per compound, from at least two independent trials per compound. The semi-adherent A2780-ADR cells exhibited a much higher variability in MTT assays, necessitating a larger sample size. For A2780-ADR cells, data are the mean ± one standard deviation with 12 samples per compound, from at least two independent trials per compound.

### Cell culture for LC-MS/MS intracellular accumulation assays

DU145-TXR cells were seeded at 350,000 cells per well in 6-well plates in complete RPMI medium. After 48 hours, the media was replaced with fresh media, and cells were treated with 5 µM of experimental compound with or without 500 nM Tariquidar (TQR), as was done previously [9]. This concentration was previously shown by us to be sufficient for inhibiting P-gp-mediated transport activity of a P-gp substrate, and was considered low enough to avoid potential interactions with the novel compounds of interest [9]. Experiments were performed in triplicate with two independent trials per compound, 6 samples, or three independent trials per compound, 9 samples total. The P-gp substrate, Daunorubicin (DAU), was included in each biological replicate as a qualitative positive control – DAU is red, so the TQR-inhibited sample with DAU turns visibly red to the naked eye. A full sample of DAU was also prepared for quantitative analysis. After a 2.5 hour incubation with the respective treatment, cells were trypsinized (0.05% Trypsin-EDTA, Gibco, ice-cold), scraped with a cell scraper, and spun at 2400 rpm. Media was removed with a glass vacuum aspirator. Cells were re-suspended and washed in 2mL of ice-cold Hank’s Balanced Salt Solution (HBSS, Gibco), and spun again at 2400 rpm. Cell lysates were diluted in 600 µL of ice-cold HBSS, flash-frozen in liquid nitrogen, and stored at -80 °C until analysis.

### Relative quantification in LC-MS/MS intracellular accumulation assays

LC-MS/MS intracellular accumulation assays were performed in collaboration with the Pharmacology Core at the University of Texas Southwestern Medical Center (UTSW). LC-MS/MS assays were performed as described in [9] with the following modification – instead of calculating a quantitative assessment of the absolute amount of compound in each sample, a relative assessment was performed. This method allowed us to quantify the change – if any – in the relative amount of compound present in the samples when TQR was added. An increase in the relative amount of compound indicates enhanced intracellular retention that compound. This quantification method can be described using the following equations:

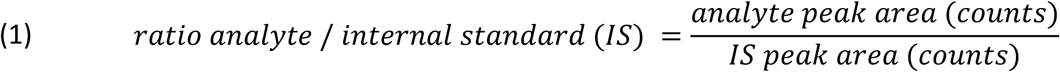

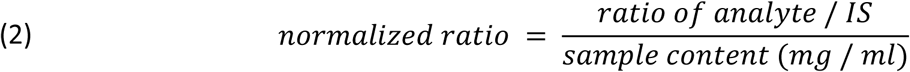

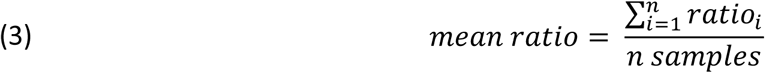

where “Analyte” refers to the experimental compound tested. The “Internal Standard” (IS) is a known substance added in a constant amount to each sample. The “Peak Area”, expressed as “counts”, describes the number of MS/MS spectra from the experimental compound and serves as a measure of its relative abundance. For each sample, the relative abundance of experimental compound is expressed as a ratio of the Analyte abundance divided by the Internal Standard abundance equation **Error! Reference source not found.**). To account for any differences in the amount of cell lysate between samples, the ratio of each sample is normalized to its respective cell lysate content expressed in mg / mL, as shown in equation **Error! Reference source not found.**). Lastly, as shown in equation **Error! Reference source not found.**), we average the “Normalized Ratio” of the samples, performed in triplicate, to produce the final values shown in S5 Table. These values allow a comparison of the relative abundance of compound between the “-TQR” and “+ TQR” samples, and to test for statistically significant differences.

### Sample preparation and analysis with LC-MS/MS

LC-MS/MS quantification was performed as follows. Cell lysate was aliquoted into Eppendorf tubes. In contrast to [9], blank cell lysates were not spiked with varying concentrations of each compound to create a standard curve, since compound amounts were normalized to abundance of the internal standard. Instead, after the initial aliquot step, MeOH containing 50 ng/mL N-Benzylbenzamide was added to each sample. Samples were vortexed briefly, incubated at 27 °C for 10 minutes, and spun at 13,200 RPM for 5 minutes. The supernatant was transferred to a new Eppendorf tube and spun once more at 13,200 RPM for 5 minutes. The supernatant was then transferred to an HPCL vial, and subsequently analyzed using LC-MS/MS using a Sciex 4000QTRAP mass spectrometer coupled to a Shimadzu Prominence LC.

Chromatography conditions were as follows: Buffer A contained water + 0.1% formic acid, and Buffer B contained methanol + 0.1% formic acid. The column flow rate was 1.5 ml/min using an Agilent C18 XDB, 5 micron packing 50 × 4.6 mm column. After the addition of the organic solvent, the remaining cell lysate was resuspended in 0.1 M NaOH, boiled for 5 min, and mixed with 1:50 B:A reagent (Thermofisher BCA Kit) to determine the cell lysate concentration in each sample. A BSA standard curve was then prepared in water and mixed using the same ratio. The samples were incubated 30 min at 37 °C and read at 562 nm. The ratio of Analyte to Internal Standard was normalized to the lysate content for each individual sample. The data for each trial are shown in S5 Table. Statistical significance was determined using a Student’s T Test in GraphPad Prism.

### Daunorubicin accumulation assays

These assays are based upon those reported in [9]. DU145-TXR cells were seeded in 96 well plates in complete RPMI medium at 15,000 cells per well. Cells were allowed to grow for 48 hours in a humidified incubator at 37 °C and 5% CO_2_. After 48 hours, media was replaced with fresh complete RPMI medium. Cells were treated with 10 µM of experimental compound with or without 10 µM daunorubicin (DAU). After a 2-hour incubation at 37 °C and 5% CO_2_ in a humidified incubator, media was removed with a vacuum aspirator, and cells were washed twice with ice-cold phosphate buffered saline (PBS) solution. Cells were lysed in PBS containing 0.5% SDS and 0.5% Triton X-100 and shaken on an Orbital Shaker for 10 min at 500 rpm. DAU fluorescence was read using the Cytation5 plate reader at excitation/emission 488 nm / 575 nm. Data are the mean ± one standard deviation from three independent experiments, 3 samples per experiment. Statistical significance was determined with a Student’s T test in GraphPad prism. None of the experimental compounds exhibited significant background fluorescence at the tested excitation and emission range.

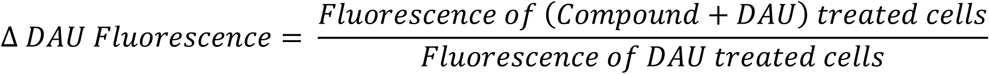

## Results

### Computational screens identify putative P-gp inhibitors for testing

We used molecular docking to computationally screen thousands of molecules for their potential binding affinity to P-gp. Docking screens were specifically designed to select for molecules that preferentially bind to the nucleotide binding domains, and that do not strongly interact with the drug binding domains. In previous work by us [7, 8], molecules were docked to a model of human P-gp that was in an closed-to-the-inside conformation [7, 20], where the nucleotide binding domains are closely engaged and the drug binding domains are widely open to the extracellular side of the membrane. Molecules that were predicted to preferentially bind the transmembrane drug binding domains were not taken into consideration for future studies.

A limitation of this earlier approach was that the closed-to-the-inside structure represents the end of the transport cycle, where pump substrates likely are released to the cell exterior. It therefore may not represent the best conformation for screening for P-gp inhibitors that are not transport substrates themselves. To identify inhibitors that are not P-gp transport substrates, it was deemed important to screen for affinity of binding to the DBD conformation at the beginning of a transport cycle, where the DBDs are open to the cytoplasm and ready to capture substrates. However, suitable open-to-the-inside structures were (and are still) not available for human P-gp. To overcome this limitation, we used targeted Molecular Dynamics (TMD) simulations to guide a model of human P-gp through conformational changes that are likely representative of a drug transport cycle [19]. These MD trajectories were then used to generate a number of open-to-the-inside and closed-to-the-inside conformations for molecular docking experiments (Fig 1). Docking search boxes were designed to sample the small molecule binding of five nucleotide binding and three drug binding domain conformations of P-gp (S1 – S2 Fig). The goal was to identify small drug like compounds that prefer the NBDs over the DBDs.

**Fig 1.**
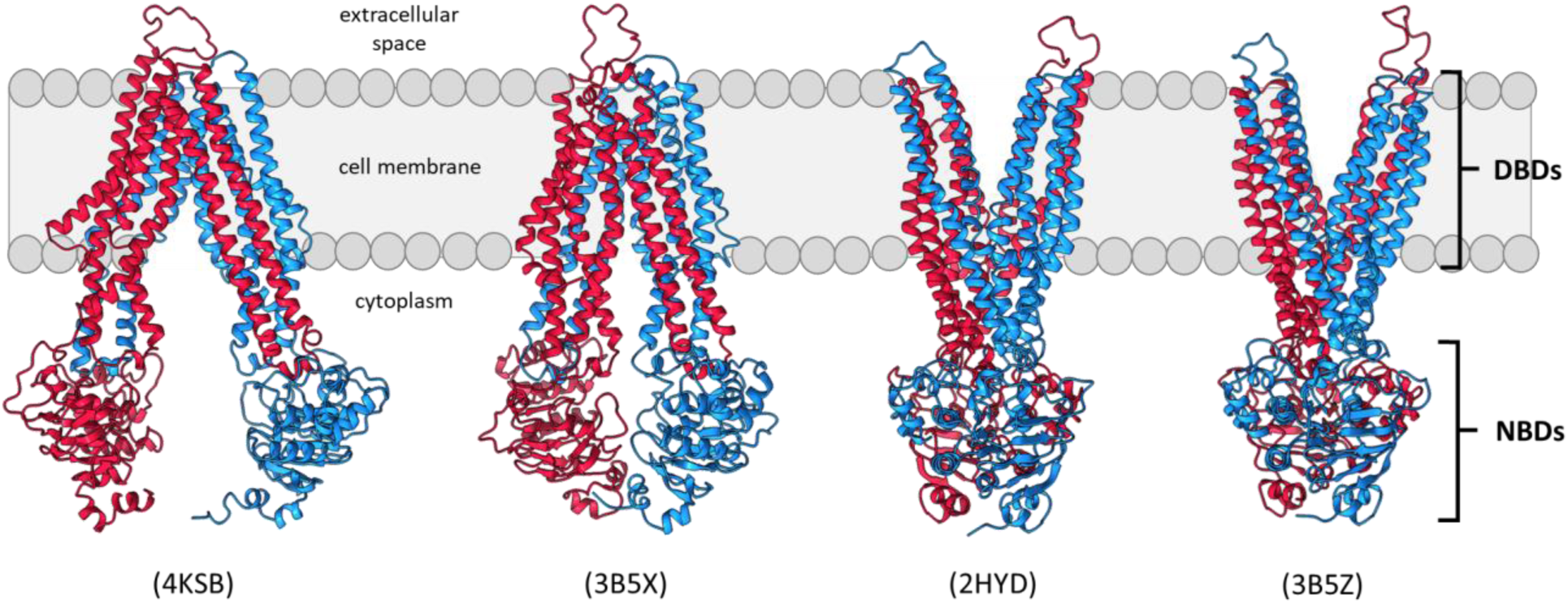
Conformations of our human P-gp model used for molecular docking screens. P-gp adopts several dynamic conformations during the substrate transport cycle. The cytoplasmic NBDs bind and hydrolyze ATP, while the membrane-embedded DBDs capture and transport substrates. P-gp switches from ‘open-to-inside/cytoplasm’ to ‘closed-to-inside’ conformation during transport, thereby alternating substrate access from the cytoplasm to the extracellular space. Search boxes for docking experiments were designed to sample the NBDs or DBDs of human P-gp in each of these conformations. The source PDB ID for each target conformation shown is in parentheses above each conformation [25, 26, 42].

AutoDock Vina [36] was used to screen molecules from the ZINC12 clean, drug-like (CDL) library against dynamic P-gp structures from TMD simulations. Our previous computational screen targeted only one conformation of P-gp; here we expand the screening to 14 different structures and locations. This increase in the number of times each molecule is iteratively screened necessitated to find a more efficient way to screen the drug libraries in an acceptable time frame. Tanimoto sets are a useful tool for reducing a molecule dataset to a more tractable size [41]. The Tanimoto algorithm determines the similarity between molecules in a dataset and uses that metric to select molecules that are representative of the chemical diversity in that dataset. In the experiments described here, Tanimoto sets were used to trim the original CDL library from ∼13 million compounds to 123,000 - 158,000 molecules (see Methods). If a molecule from the Tanimoto set performed well in docking screens, similar molecules were collected from the CDL dataset and screened against the same P-gp targets.

Using this iterative approach, 95% of the chemical diversity within the CDL library was efficiently screened for estimated affinity to the DBDs and the NBDs of P-gp. Molecules with <u>low</u> estimated affinities at the <u>DBDs</u> and *high* estimated affinities at the *NBDs* were retained during each iterative round of molecular docking, as described in [7]. After the final docking screen, the top 100 molecules against each P-gp target structure were subjected to quantitative structure-activity relationship (QSAR) predictions using the Online chemical database (OCHEM) [37]. The QSAR data, select predicted chemical properties, and estimated binding affinities were used to select 67 compounds for testing with cell-based assays. The experimental compounds were numbered 58 to 124 in order of arrival to the laboratory.

### Potential P-gp inhibitors reverse MDR in P-gp-overexpressing prostate cancer cells

The computationally identified compounds were screened for the ability to re-sensitize P-gp overexpressing DU145-TXR cancer cells to paclitaxel (PTX), a chemotherapeutic drug and transport substrate of P-gp [38]. DU145-TXR cells greatly overexpress P-gp relative to the DU145 parental cells and exhibit a 34-fold increase in the IC_50_ of PTX relative to that of DU145 cells [38]. In the described experiments, re-sensitization was defined as a 30% or greater decrease in cell viability between cells treated with PTX alone, or with a combination of PTX and the experimental compound. DU145-TXR cells were exposed to compounds 58 to 124 at 15 µM, with or without 500 nM PTX, for 48 hours. Cell viability was then assessed with MTT viability assays as previously described [9]. Tariquidar and verapamil were included as positive controls for P-gp inhibition, and the BCRP inhibitor Ko143 was included as a negative control for P-gp inhibition [18]. Treatment with 11 compounds (61, 68, 70, 78, 96, 97, 101, 103, 111, 122, 124) re-sensitized DU145-TXR cells to PTX (Fig 2A, S4 Table). The remaining compounds were eliminated from further consideration as potential P-gp inhibitors.

**Fig 2.**
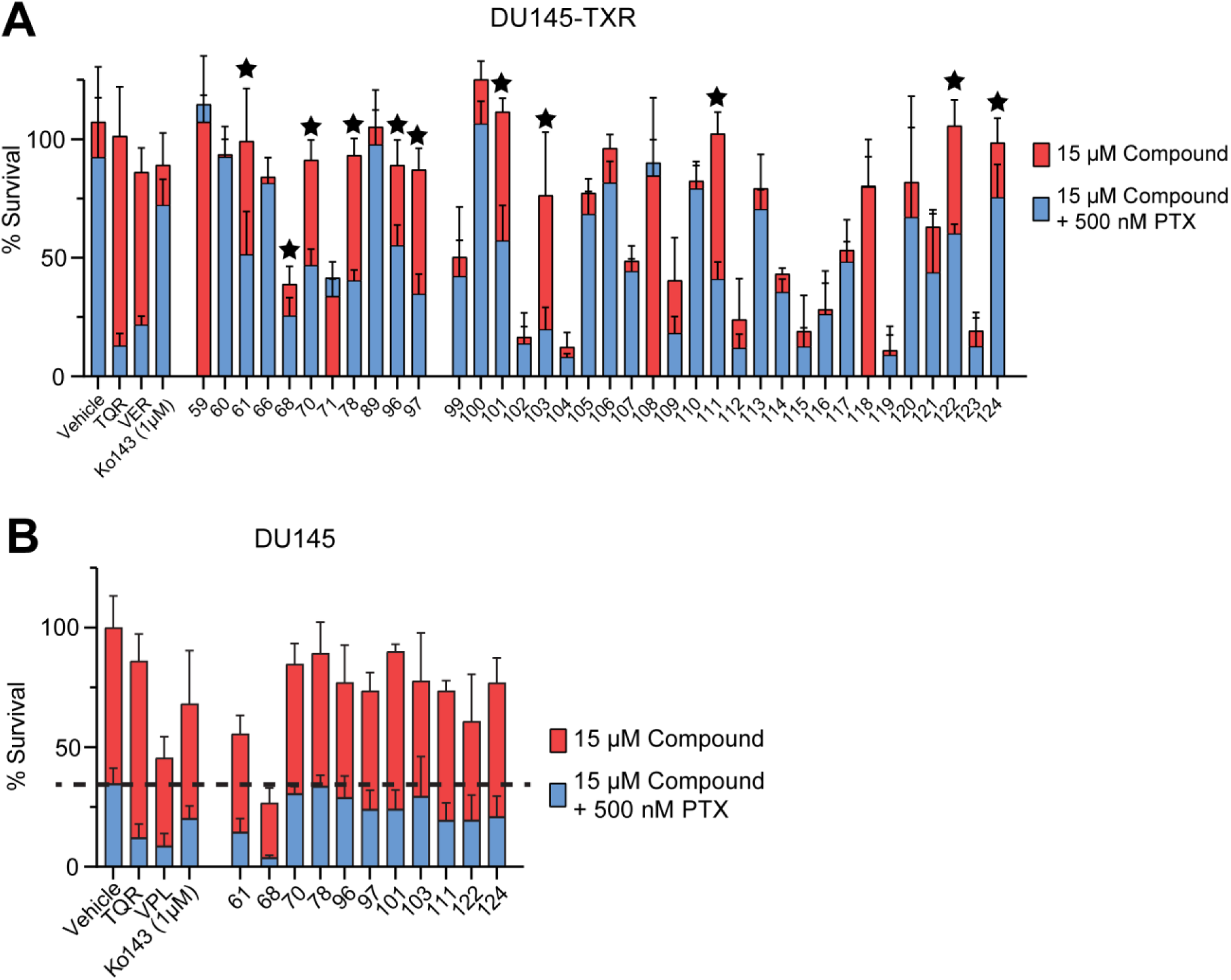
Potential P-gp inhibitors reverse MDR in P-gp overexpressing DU145-TXR cancer cells. 67 compounds were chosen from computational screens for testing *in vitro*. **A)** DU145-TXR cells were exposed to 15 µM experimental compound with or without 500 nM of the P-gp substrate and chemotherapeutic paclitaxel (PTX). Tariquidar (TQR) and verapamil (VPL) at 15 µM were used as positive controls for P-gp inhibition, while Ko143 at 1 µM and DMSO vehicle control were used as negative controls for P-gp inhibition. Percent (%) survival is calculated as the percent survival relative to that of the vehicle control (DMSO-treated) cells (see Methods). Starred compounds are considered ‘top hits’. **B)** Top compounds were tested for inherent toxicity with the non-P-gp overexpressing DU145 parental line. Cells were exposed to 15 μM compound with or without 500 nM PTX for 48 hours, after which cell viability was assessed using the MTT viability assay. Data represent the mean ± one standard deviation from the mean (n = 8, two independent trials).

The 11 compounds that re-sensitized DU145-TXR cells to PTX (Fig 2A) were then tested for their ability to sensitize the parental line DU145 cells to PTX [38]. Our hypothesis was that since DU145 cells do not overexpress P-gp, a compound that enhances PTX toxicity in DU145 cells is unlikely to act solely by inhibiting P-gp transport, but may have a different cellular target. In these experiments, DU145 cells were exposed to the experimental compounds at 15 µM in the presence or absence of PTX and cell viability was assessed using MTT assays. Nine of the 11 compounds tested (compounds 70, 78, 96, 97, 101, 103, 111, 122, and 124) were observed to insignificantly sensitize DU145 cells to PTX (Fig 2B) as compared to exposing the cells to PTX alone, see dotted line and PTX toxicity in the absence of compounds. Except for compound 68, the top hits were not observed to exhibit significant intrinsic toxicity against DU145 cells. Verapamil is known to cause significant off-target effects [11], so its effect on non-P-gp overexpressing DU145 was not really surprising.

### Potential P-gp inhibitors are not specific to one cancer cell line

To test if the potential P-gp inhibitors were specific to DU145-TXR cancer cells, the top compounds (70, 78, 96, 97, 101, 103, 111, 122 and 124) were tested with second set of paired cancer cell lines – the P-gp overexpressing, doxorubicin resistant A2780-ADR ovarian cancer cell line, and the parental, non-P-gp overexpressing A2780 line [9, 43, 44]. Doxorubicin is both a chemotherapeutic and a P-gp transport substrate. Since it is possible that the resistance of the A2780-ADR cell line to Doxorubicin involves mechanisms other than P-gp overexpression, assays used PTX to test for the potential to reverse MDR. Tariquidar and verapamil were used as positive controls for P-gp inhibition, with the BCRP inhibitor, Ko143, as a negative control. Compound 59 was used as an additional negative control because it did not reverse MDR in DU145-TXR cells (Fig 2A). Of the nine compounds tested against DU145-TXR, each compound (70, 78, 96, 97, 101, 103, 111, 122, and 124) also reversed MDR in A2780-ADR cells (Fig 3A), but did not greatly increase the toxicity of PTX against the parental A2780 cells (Fig 3B, dashed line represents the effect of 500 nM PTX alone). These nine compounds (70, 78, 96, 97, 101, 103, 111, 122, and 124) were therefore considered to be good candidates for further toxicity testing and for evaluation as potential P-gp pump substrates.

**Fig 3.**
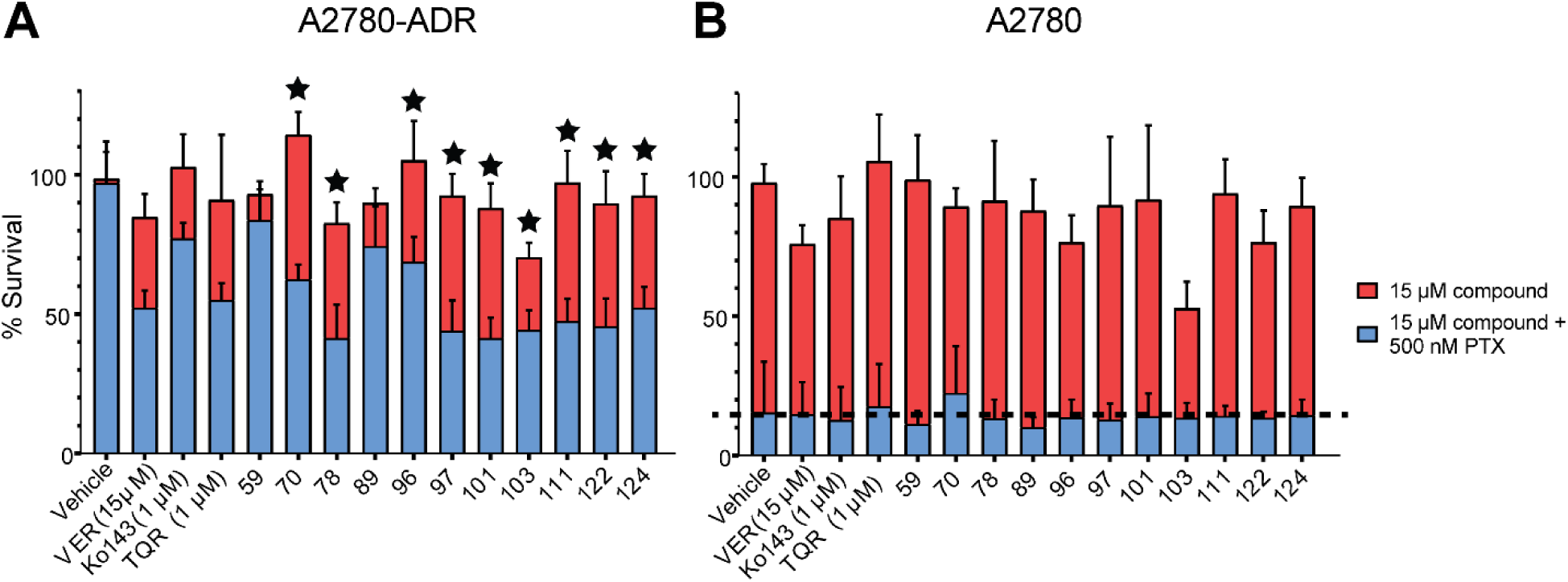
Putative P-gp inhibitors reverse MDR in P-gp-overexpressing A2780-ADR cancer cells. The top 10 compounds from screens with DU145 and DU145-TXR were tested at 15 μM concentration against **A)** the chemotherapy-resistant, P-gp overexpressing A2780-ADR line and **B)** the chemotherapy-sensitive, non-P-gp overexpressing parental A2780 line. Cells were exposed to compounds with or without 500 nM Paclitaxel (PTX) for 48 hours. The vehicle (DMSO) control represents PTX toxicity in the absence of compound (dotted line). Cell viability was assessed using the MTT viability assay. Data represent the mean ± one standard deviation from the mean, 8 replicates per compound, from at least two independent trials. Compounds 59 and 89 were used as internal negative controls for compounds that did not reverse MDR in DU145-TXR cells (Fig 2A). Starred compounds in panel **A)** were considered to be the top hits.

### Compounds are not substantially toxic to non-cancerous human cells

If a P-gp inhibitor is intrinsically toxic to non-cancerous cells, it is more likely to be participating in undesirable off-target interactions. The top compounds from the screens shown above were screened for inherent toxicity using the non-cancerous HFL1 (human lung fibroblast) cell line [45]. HFL1 cells were incubated with 15 µM compound for 48 hours, after which cell viability was assessed using MTT cell viability assays. A 20% reduction in cell viability (compared to DMSO-treated cells) was considered the maximum acceptable toxicity (see dotted line). The chemotherapeutic daunorubicin was included as a positive control for toxicity. Nine compounds (70, 78, 96, 97, 101, 103, 111, 122 and 124) were deemed by us as either ‘non-toxic’ or ‘within acceptable limits’ of toxicity (Fig 4).

**Fig 4.**
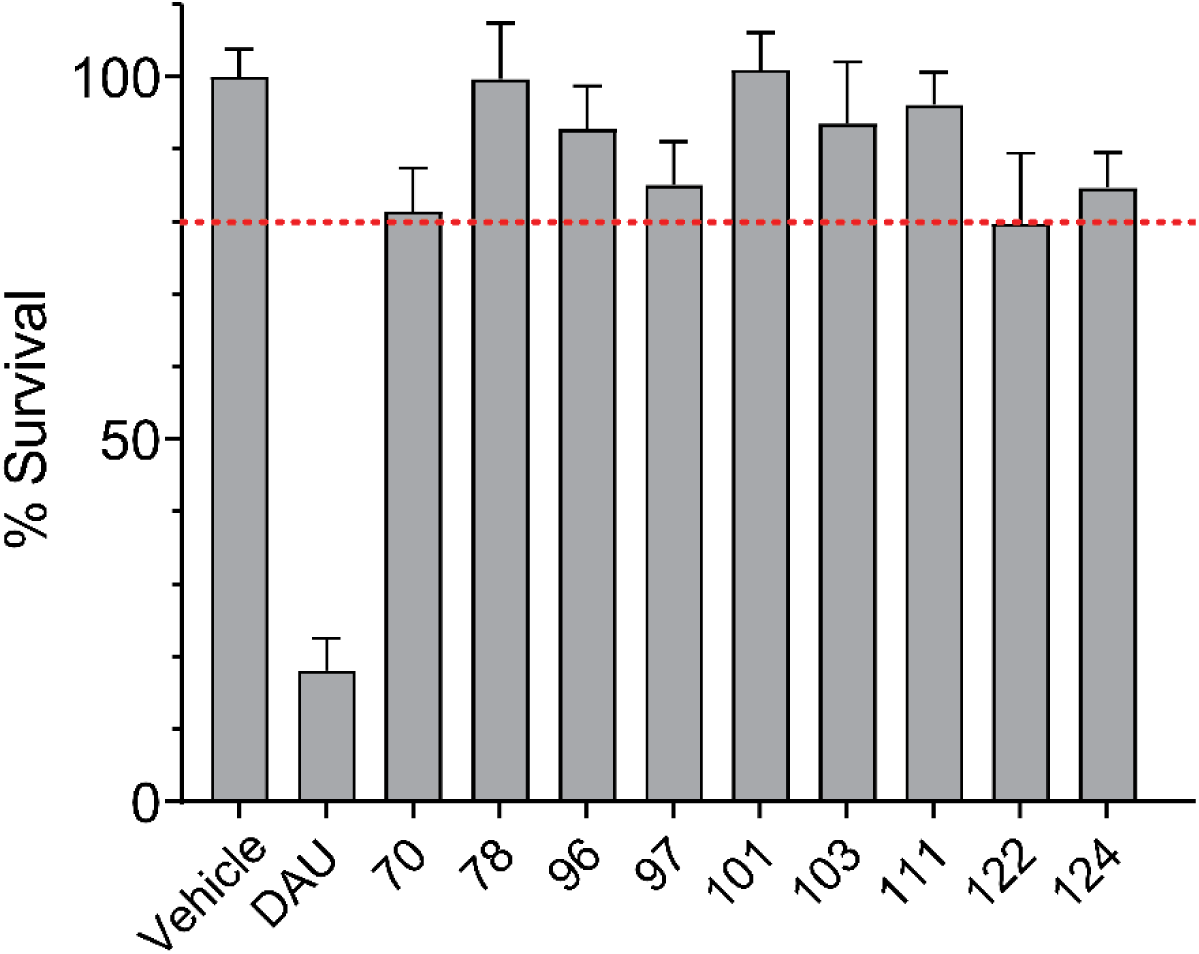
Evaluating the inherent toxicity of potential P-gp inhibitors. The top compounds from previous screens were tested against the non-cancerous HFL1 cell line. HFL1 cells were exposed to 15 µM of experimental compound. Daunorubicin (DAU) was used as a positive control for inherent toxicity. Cells were exposed to the experimental treatments for 48 hours, after which cell viability was assessed using MTT viability assays. Data represent the mean ± one standard deviation from the mean, 8 samples per compound, from at least two independent experiments. The dashed line marks 80% viability of DMSO-treated cells. All of the experimental compounds were deemed to have tolerable toxicity.

### Potential P-gp inhibitors are not transport substrates of P-gp

We aimed to identify P-gp inhibitors that are not transport substrates of the pump itself. To that end, our subtractive docking methods selected molecules that were predicted to favor the NDBs over the DBDs. To test these predictions *in vitro*, liquid chromatography with tandem mass spectrometry (LC-MS/MS) was used to assess the intracellular accumulation of experimental compounds as described in [9]. The P-gp overexpressing DU145-TXR cells were exposed to 5 µM compound with or without 500 nM tariquidar for 2.5 hours. At 500 nM concentration, tariquidar should strongly inhibit P-gp while not being transported itself [16]. The relative intracellular accumulation of each experimental compound was then assessed using LC-MS/MS. If the compound is a P-gp transport substrates, it should accumulate intracellularly in the presence of tariquidar (P-gp inhibited), while the intracellular concentration should be lower in the absence of tariquidar (P-gp active). The P-gp substrate and chemotherapeutic daunorubicin (DAU) was used as a positive control for P-gp-mediated transport. Six of the compounds that reversed MDR in DU145-TXR and A2780-ADR cells (70, 78, 96, 97, 101, and 111) were deemed unlikely to be transport substrates of P-gp (Table 1 and S5 Table). Three compounds (103, 122 and 124) were found to be potential P-gp transport substrates.

**Table 1.**
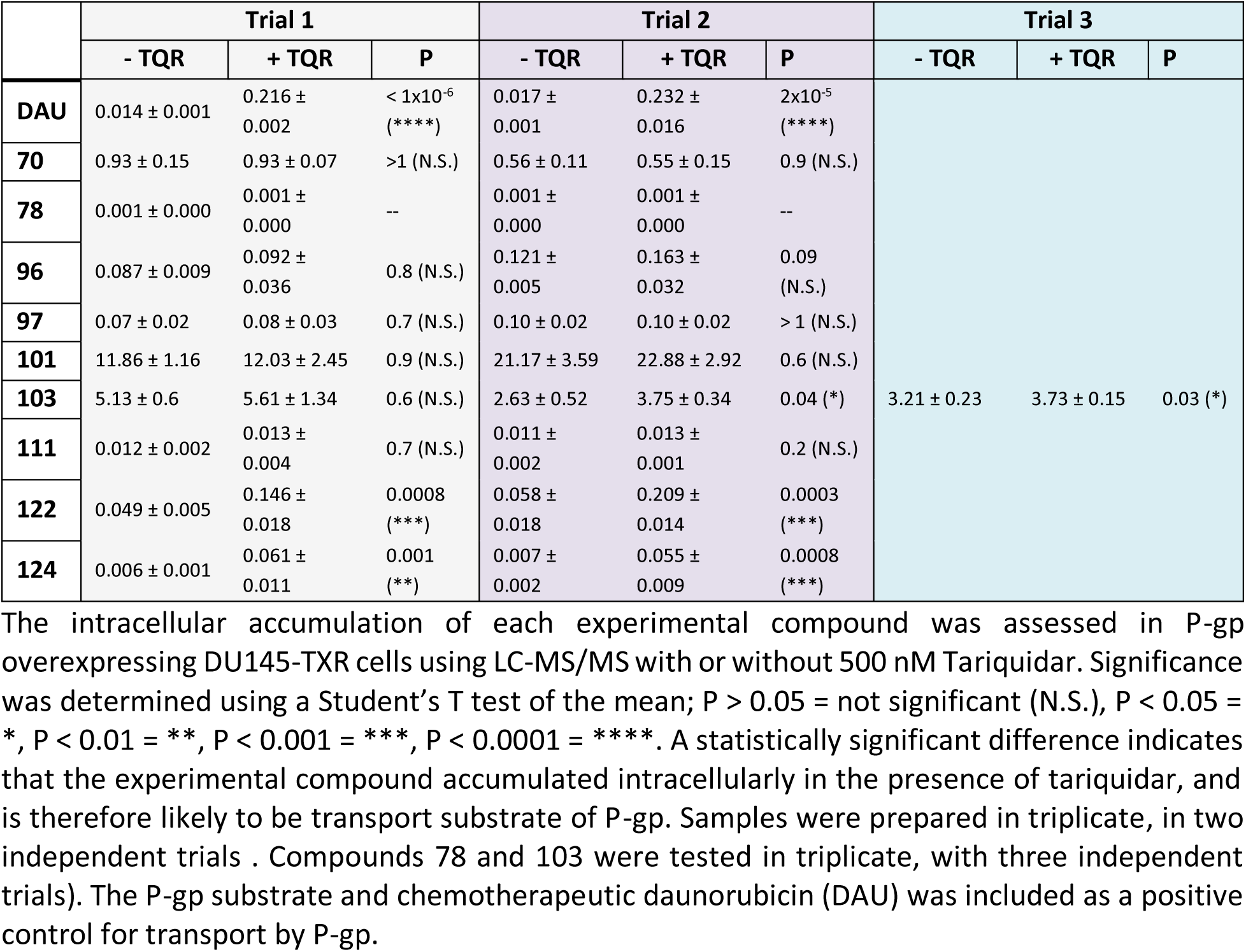
Intracellular accumulation of compounds measured by LC-MS/MS.

**Table 2.**
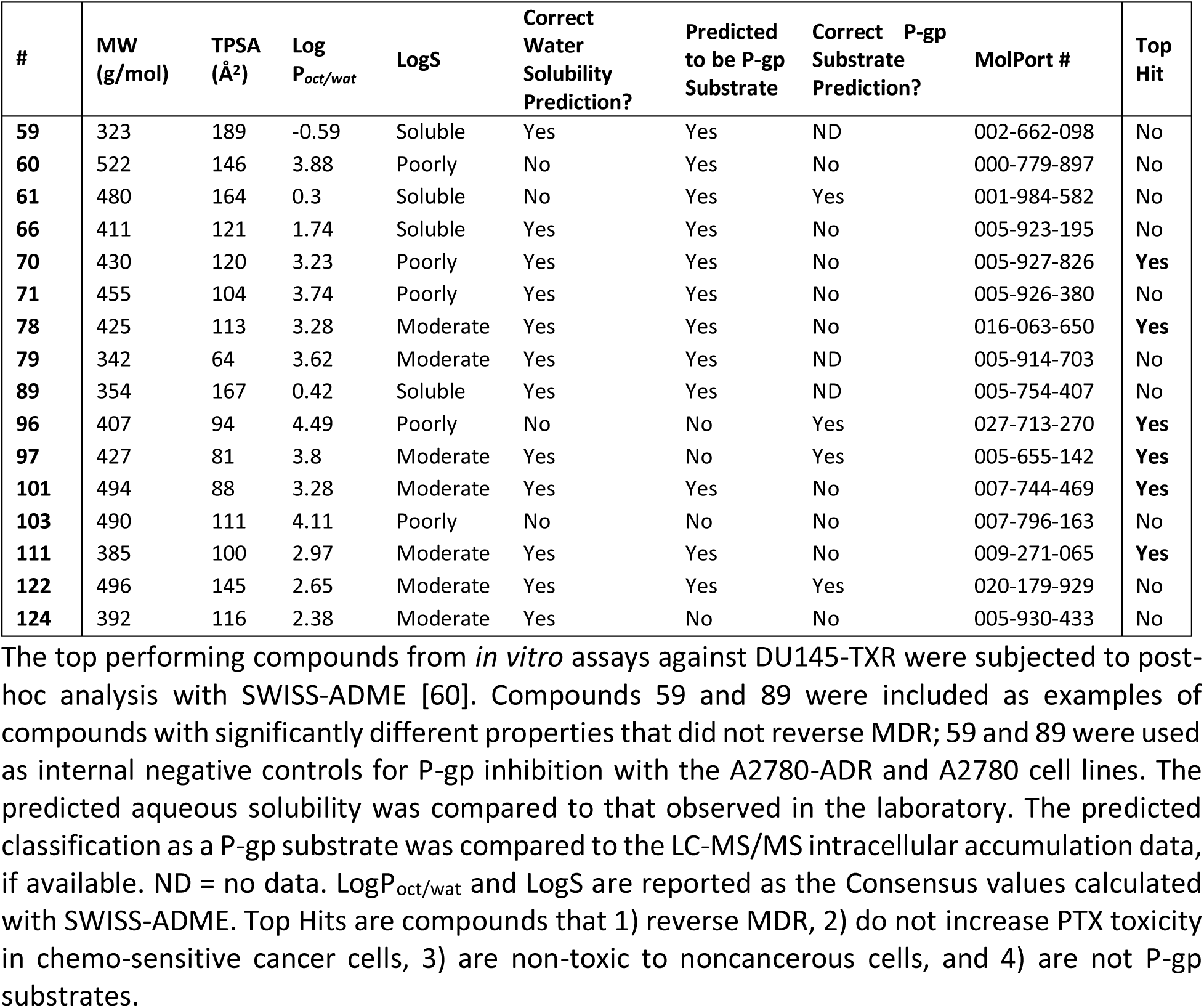
Computational analysis of compounds and comparison to *in vitro* data.

### Experimental compounds inhibit the transport of a P-gp substrate

To ensure that the top candidates from our previous studies were indeed targeting and inhibiting P-gp activity, we tested these compounds for their ability to enhance intracellular accumulation of the P-gp substrate, daunorubicin (DAU). These assays used the P-gp highly overexpressing DU145-TXR cell line. In this assay, the intrinsic fluorescence of DAU was used to quantify its relative concentration in the cell. An increase in fluorescence correlates with an increase in the intracellular accumulation of DAU. Compounds 70, 78, 96, 97, 101, 103, 111 were assessed for the ability to increase the intracellular retention of DAU. Compound 59 was included as a negative control for P-gp inhibition, tariquidar and verapamil were included as positive controls for P-gp inhibition. Cells were treated with 10 µM of experimental compound with or without 10 µM DAU. After a 2-hour incubation, cells were washed and lysed, and the total fluorescence of DAU in each well was quantified. Treatment with compounds 70, 96, 97, and 101, and 103 significantly increased the DAU fluorescence in DU145-TXR cells, indicating an increase in the intracellular retention of DAU (Fig 5, S4 Fig, S6 Table). As expected, treatment with compound 59 did not significantly increase fluorescence of DAU. The results of these assays suggest that compounds 70, 96, 97, 101, and 103 significantly inhibited transport of DAU by P-gp. Compounds 78 and 111 did not cause an increase in intracellular DAU retention. Together with LC-MS/MS data, our results suggest that compounds 70, 96, 97, and 101 are not P-gp substrates, and therefore are more likely to bind at the cytoplasmic NBDs, which were targeted our docking studies.

**Fig 5.**
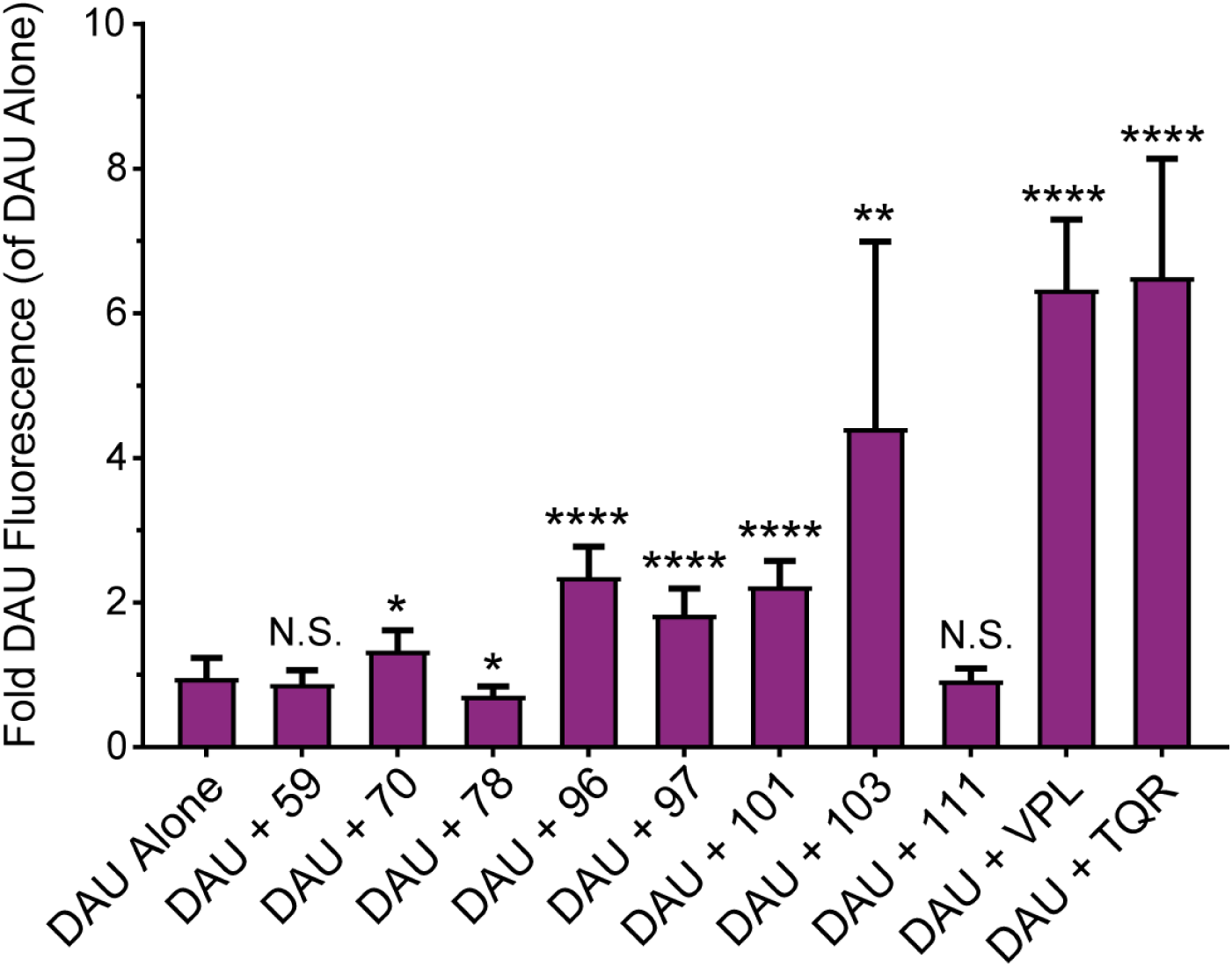
Change in Fluorescence of DAU in the Presence of Experimental Compounds. Fold change in fluorescence of the P-gp substrate Daunorubicin (DAU) in the presence or absence of experimental compounds or known P-gp modulators. Cells were treated with 10 µM compound in the presence or absence of 10 µM DAU. After washing and lysing the cells, the intracellular DAU fluorescence was measured and expressed as a fold change relative to the fluorescence of DU145-TXR cells treated with DAU alone. The P-gp inhibitors, VPL and TQR, were included as positive controls for P-gp inhibition, and compound 59 was included as a negative control for P-gp inhibition. Three samples per trial, three independent trials. Significance was determined using a Student’s T test of the mean by comparing fluorescence of DAU and compound to that of DAU alone; P > 0.05 = N.S., P < 0.05 = *, P < 0.01 = **, P < 0.001 = ***, P < 0.0001 = ****. Data also shown in S4 Fig.

## Discussion

### The search for P-gp inhibitors to treat multidrug resistant cancer

Multidrug resistance is a major obstacle to the treatment of human cancer. As prominent agents of multidrug resistance, ABC transporters like P-gp are clinically-relevant targets and the focus of decades of study [2]. However, P-gp has proven notoriously difficult to target in drug discovery, and the pump’s dynamic conformational changes lie at the heart of the difficulty. During transport, P-gp undergoes substantial conformational changes that significantly rearrange the nucleotide binding domains (NBDs) and drug binding domains (DBDs) (Fig 1). These conformational changes cause the ‘topology’ or structural landscape of P-gp – and thus, the binding sites for potential inhibitors – to change dynamically during transport. Adding to the difficulty, knowledge of P-gp’s transport cycle remains limited because it is not possible to observe the conformational changes of the protein directly. Recent cryo-EM work has provided unprecedented insight into the conformational changes of P-gp [5, 16], but these structures were not available when our docking studies and simulations were performed. Our drug discovery pipeline was successful regardless, and we expect that future efforts will only be enhanced by the availability of higher-resolution structures.

### The search for P-gp inhibitors has historically targeted the drug binding domains

Efforts to target and inhibit P-gp have historically focused on the pump’s transmembrane drug binding domains (DBDs). The DBDs are hydrophobic, flexible, and large enough to accommodate molecules as bulky as the Alzheimer’s associated amyloid β peptides (∼4000 Daltons)[6]. In prior work, we also showed that the DBDs are large enough to allow substrates to take multiple paths through the protein during transport [19], indicating that binding sites for small molecule inhibitors in the DBDs are both abundant and variable. Indeed, some of the best-characterized P-gp inhibitors (see S1 Table), including Tariquidar and Verapamil, bind P-gp at the DBDs [16, 46]. Interestingly, Tariquidar and Verapamil have been shown to be P-gp transport substrates, and both compounds failed clinical trials due to off-target toxicity exacerbated by high therapeutic doses. Drug discovery screens targeting P-gp are fraught with failure, leading to alternative approaches such as the testing of natural compounds [47–49] as P-gp inhibitors. Despite the difficulty in designing effective inhibitors, P-gp remains an important target in the fight against multidrug resistance, and an active subject of drug discovery screens [2, 50–52].

For P-gp inhibitors, poor performance in clinical trials appears to be correlated with binding at the DBDs, and most notably, with being a transport substrate of the pump itself (S1 Table). If an inhibitor is a transport substrate of P-gp, the required therapeutic dose is higher than for a non-substrate because the pump actively effluxes its own inhibitor from the cell. We speculate that the combination of these two factors – preferential binding to the DBDs, and being transported by P-gp – are a major factor in why previous generations of P-gp inhibitors have failed in clinical trials.

### The nucleotide binding domains of P-gp are an underappreciated target

In earlier work we demonstrated that the pump’s cytoplasmic NBDs are viable targets for drug discovery [7, 8]. Our first-draft attempt used molecular docking to screen for molecules that were predicted to prefer the NBDs over the DBDs. Even though this study used a homology model of human P-gp, and only one conformation of the pump, the pipeline successfully identified four compounds that reversed MDR in cancer cells, with an impressive ∼ 7% hit rate [7, 8]. Subsequent work by us showed that chemical and structural optimization of one compound, SMU 29, yielded more even novel P-gp inhibitors that reversed MDR in cancer cells [10, 53]. In addition to our work, a recent study used a similar technique to identify more novel inhibitors that target the NBDs [54]. Taken together, these results demonstrate that the NBDs of P-gp are viable targets for drug discovery.

### Our enhanced, computationally-assisted drug discovery pipeline

Here we aimed to identify novel, chemically-diverse P-gp inhibitors that reverse MDR in human cancers. To do so, we built upon the success of our earlier work and designed an enhanced computationally-accelerated pipeline with the following features. First, our original pipeline used a model of human P-gp in only one conformation. To better capture the conformational complexity of P-gp during transport, we used TMD simulations with our P-gp model to generate a suite of conformations for molecular docking. Second, the original screen used a limited molecule dataset. Here we iteratively screened millions of compounds against the NBDs and DBDs of *each* dynamic structure, using Tanimoto sets to enhance computational efficiency while maximizing the chemical diversity of screened ligands. Third, our original approach used one pair of resistant and non-resistant cancer cell lines and did not consider toxicity to non-cancerous cells. Here we tested compounds against two pairs of resistant/non-resistant human cancer cell lines, and screened for inherent toxicity with a non-cancerous human cell line. Finally, we incorporated assays to test whether our compounds are likely to be transport substrates of P-gp. Using this combination of *in silico* and *in vitro* techniques, the pipeline presented here aims to identify putative inhibitors of P-gp that preferentially target the NBDs over the DBDs.

### Success of the pipeline in identifying novel inhibitors of P-gp

The validity of our computationally-accelerated approach is demonstrated by its success: of the 67 compounds identified through docking, nine reversed MDR in human cancer cell lines, a 13.4% hit rate for compounds that reverse MDR. These compounds were shown to be non-toxic to non-cancerous cells, and all were effective in the 15 µM range. Of the nine compounds, six were shown to be unlikely transport substrates of P-gp. Each compound is a good candidate for lead optimization and testing (Fig 6).

**Fig 6.**
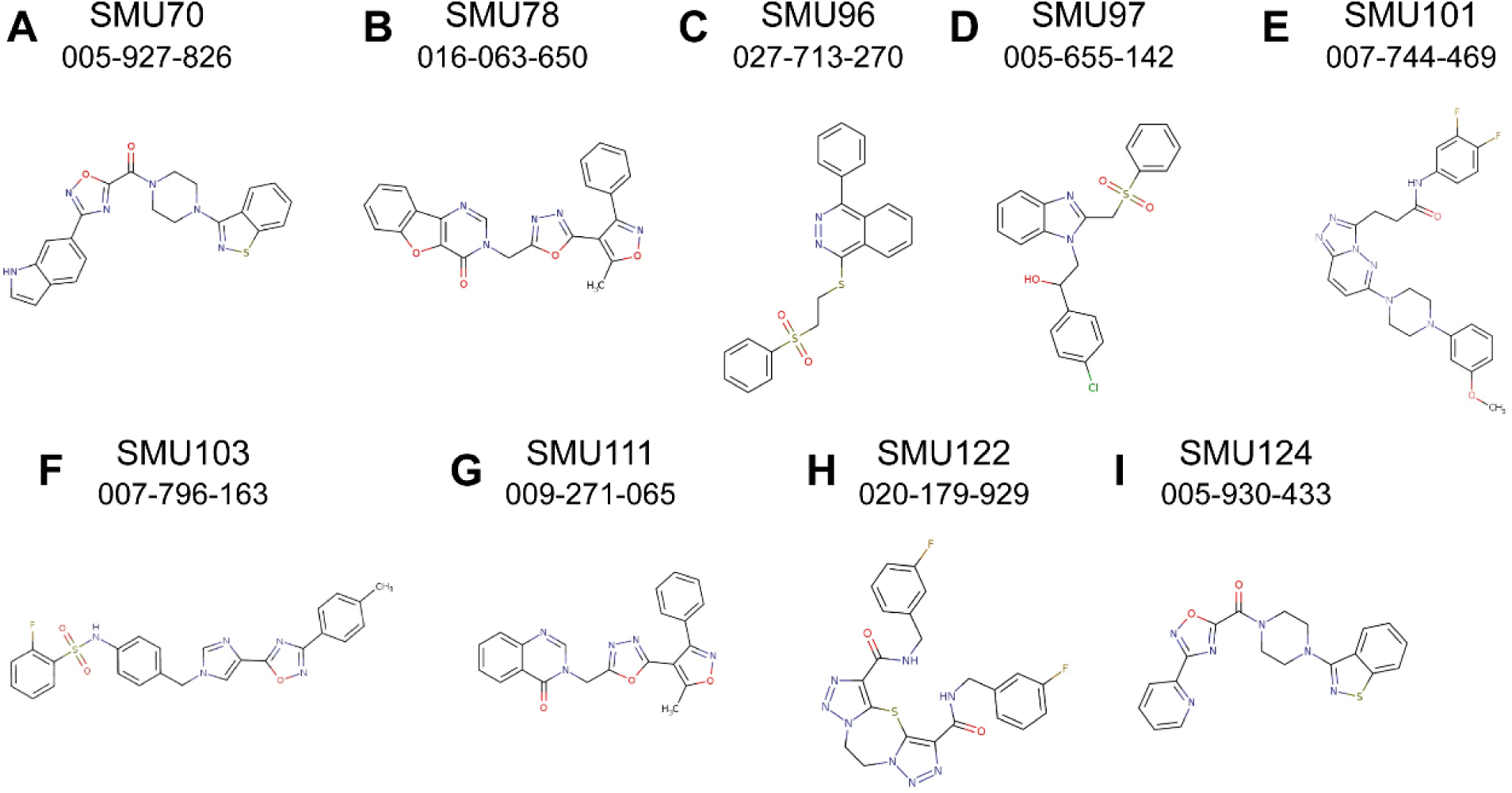
Structures of novel P-gp inhibitors identified by virtual-assisted drug discovery. Chemical structures of the nine best P-gp inhibitors identified in this study (panels A through I) with our laboratory name (SMU-) above, and the MolPort identification number below. The ZINC identification number, IUPAC International Chemical Identifier (InChI) key, and the Simplified Molecular Input Line Entry System (SMILES) code for each molecule are listed in S7 Table.

### Comparison to other computationally-assisted screens targeting P-gp

Our pipeline has a 13.4% success rate for compounds that reverse multidrug resistance, but importantly, these compounds are effective at 15 µM. Combined with the 13.4% hit rate, an effective dose of 15 µM represents an enhancement of traditional computational screening methods by a significant factor. For identification of lead compounds that are effective at sub–100 μM concentrations (6-fold greater than 15 µM reported here), the median hit rate of virtual-assisted efforts is approximately 13% [55]. 103 studies reported hit rates greater than 25%; however, 75% of these studies tested fewer than 20 compounds and used a sub-100 μM dose as the minimal dose cutoff [56]. These trends of small sample size and very high dose concentration cut-offs have been observed in other virtual-assisted studies [57–59]. In a recent computationally-assisted study targeting the P-gp NBDs, the effective dose of novel compounds ranged from ∼300 to 1033 µM [54].

### Success of pipeline despite lack of high-resolution structures of P-gp

Many virtual-assisted studies benefited from high-resolution structures of the target proteins, frequently including co-crystallized inhibitors. It is crucial to note that, when the docking studies presented here were performed, high resolution crystal or cryo-EM structures of human P-glycoprotein at multiple stages of transport were not available. Even to date, a complete suite of structures encompassing the full catalytic transport cycle of human P-glycoprotein is not available. As an additional factor, when our docking studies were performed, the available structures of P-gp homologues were limited in both resolution and conformational diversity. To overcome this limitation and produce suitable structures for molecular docking, we built a homology model of human P-gp and used targeted MD simulations to generate structural ‘snapshots’ of the transporter throughout a putative catalytic drug transport cycle [19, 20]. Our first-draft virtual screening pipeline with our P-gp homology model resulted in a 7% hit rate for compounds that reverse multidrug resistance in P-gp-overexpressing cancer cells [7, 8]. We also provided evidence that these small molecules inhibited substrate-stimulated ATPase activity of P-gp, suggesting interactions with the NBDs – the overall goal of the docking screens [7]. Despite the limited access to conformationally diverse and high-resolution structures of human P-gp when our docking studies were performed, the work presented here shows a two-fold increase (13.4%) in hit rate for compounds that reverse multidrug resistance in P-gp-overexpressing cancer cells, thereby validating the use of our P-gp model to virtually screen small molecules with molecular docking, and the use of counter-selective docking to target the NBDs.

### Validation and testing of the SWISS-ADME computational chemical analysis

Identifying P-gp inhibitors that are not transport substrates remains inherently challenging. To that end, computational tools such as the SWISS-ADME server attempt to use a molecule’s unique chemical and structural characteristics to predict whether it is likely to be a P-gp transport substrate [60]. Our LC-MS/MS assays allowed us to retroactively test the validity of predictions from SWISS-ADME. Of 13 compounds assayed in total using LC-MS/MS (not all data shown), only four were correctly predicted to be P-gp transport substrates. This discrepancy underscores the inherent difficulty of computationally predicting transport substrates of this dynamic ABC transporter. We have also included the SWISS-ADME analyses of molecular weight, topological polar surface area, Log P_oct/wat_ (hydrophobicity), and Log S (water solubility) for our top hits from this work, as well as for several compounds that did not reverse MDR, in hopes that these data can be used to guide selection of future P-gp inhibitors for screening. Alternative identifiers for the molecules identified in this study are shown in S7 Table.

### Adapting this pipeline to target other ABC transporters

The ABC transporter superfamily contains many proteins that undergo similar conformational dynamics to P-gp [2, 3]. One of the most clinically-relevant is breast cancer resistance protein (BCRP, *ABCG2*), which confers MDR to several types of human cancers [61–63]. Similar to P-gp, BCRP also consists of cytoplasmic NBDs and membrane-embedded DBDs, and previous drug discovery efforts have focused on the DBDs [64]. DBD-targeted BCRP inhibitors have exhibited problems with poor solubility, principally because molecules that bind to the DBDs tend to be hydrophobic [64]. The NDBs of BCRP have not been thoroughly explored in drug discovery, and our results suggest that the NBDs are good candidates for virtual-assisted screening and the pipeline presented here. High-resolution structures of BCRP are increasingly available in several conformations, which will enhance drug discovery efforts [65–68]. As an additional factor, paired resistant/non-resistant, BCRP-overexpressing human cancer cell lines are available, which will facilitate *in vitro* testing of compounds [9].

In addition to conferring MDR to cancers, ABC transporters like MsbA from *E. coli* also perform essential functions in pathogenic bacteria [42, 69, 70]. Targeting bacterial ABC transporters like MsbA with the NBD-targeting docking methods presented here could identify novel drug candidates to aid in the fight against drug resistant bacterial infections [69].

The nucleotide-binding region of ABC transporters contains several conserved residues, specifically those that are involved in ATP binding and hydrolysis [42]. While this conservation might cause an NBD-targeting P-gp inhibitor to bind other ABC transporters, one might argue that this ‘off-target’ activity against other ABC transporters is acceptable within the context of multidrug resistant cancer – for these patients, chemotherapy has failed, options are limited, and more than one ABC transporter is potentially at play in conferring multidrug resistance. P-gp is not the only ABC transporter that confers multidrug resistance [62, 71], and the chemotherapy substrate profiles of ABC transporters often overlap. Therefore, it is possible that targeting several ABC transporters could be effective within the context of drug resistant cancer. This idea remains to be tested, as there are currently no NBD inhibitors available to patients, but future efforts should consider activity against other ABC transporters.

In summary, the methodology presented here, particularly the use of counter-selective docking methods that target the NBDs over the DBDs, can be readily adapted to target other members of the ABC transporter family. Following recent advances in cryo-EM, high-resolution structures of ABC transporters are increasingly available for use in drug discovery screens [5, 16]. These high-resolution structures were not available when the simulations and docking for this study were performed. Our computationally-assisted pipeline was successful despite the lack of high-resolution structures, which underscores the untapped potential of our approach future drug discovery screens targeting P-gp (and other ABC transporters) will benefit immensely from the growing body of high-resolution structures, particularly those derived by cryo-EM. The nucleotide-binding regions of ABC transporters are

## Supporting information

S8_Cell_Culture_Data

S9_Docking_Data

## Acknowledgements

We would like to thank Ms. Jesiska Lowe for her help in preparing the LC/MS-MS accumulation assays.

## Author’s contributions

Lauren A. McCormick – conceptualization, visualization, methodology, analysis, writing – original draft, review, editing

James W. McCormick – conceptualization, visualization, methodology, analysis, writing – original draft, review, editing

Chanyang Park – investigation, visualization, editing

Courtney A. Follit – investigation, conceptualization

John G. Wise – conceptualization, supervision, funding acquisition, review and editing

Pia D. Vogel – conceptualization, supervision, funding acquisition, review and editing

## Consent for Publication

The authors have given their consent to publish.

## Copyright

© The Authors 2024

## Supporting information

### Supplemental figures and tables

**S1 Fig.**
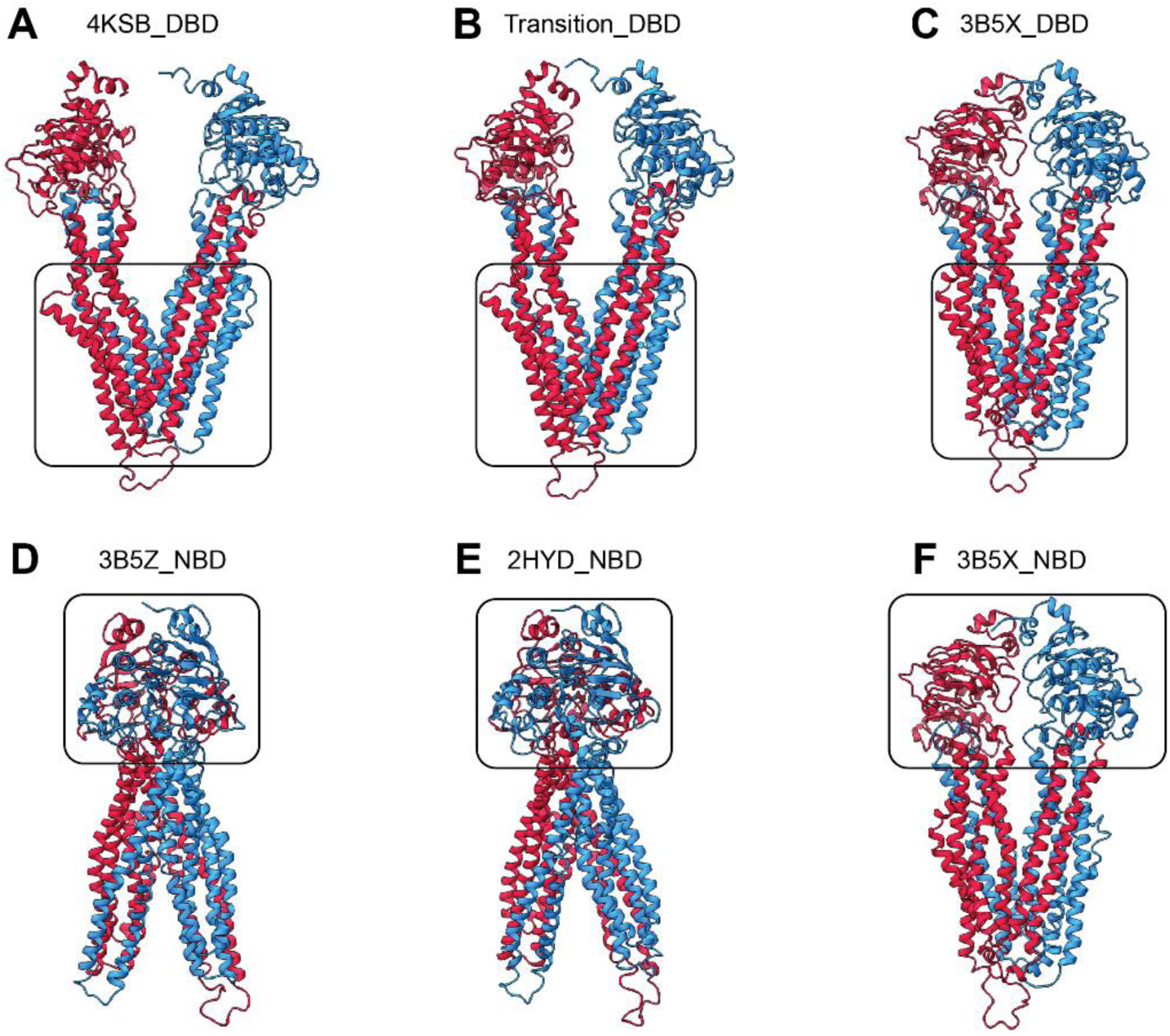
**Target structures for docking studies**. **Panel 1:** A. 4KSB_DBD, B. Transition_DBD, C. 3B5X_DBD, D. 3B5Z_NBD_1 and 3B5Z_NBD_2, E. 2HYD_NBD_1 and 2HYD_NBD_2, F. 3B5X_NBD. The NBD of the Transition structure in B was also sampled. NBD dock boxes for the 2HYD and 3B5Z structures were designed to sample each NBD individually. A-F are human P-gp structures. Boxes correspond to the region targeted using AutoDock Vina with an exhaustiveness of 128.

**S2 Fig.**
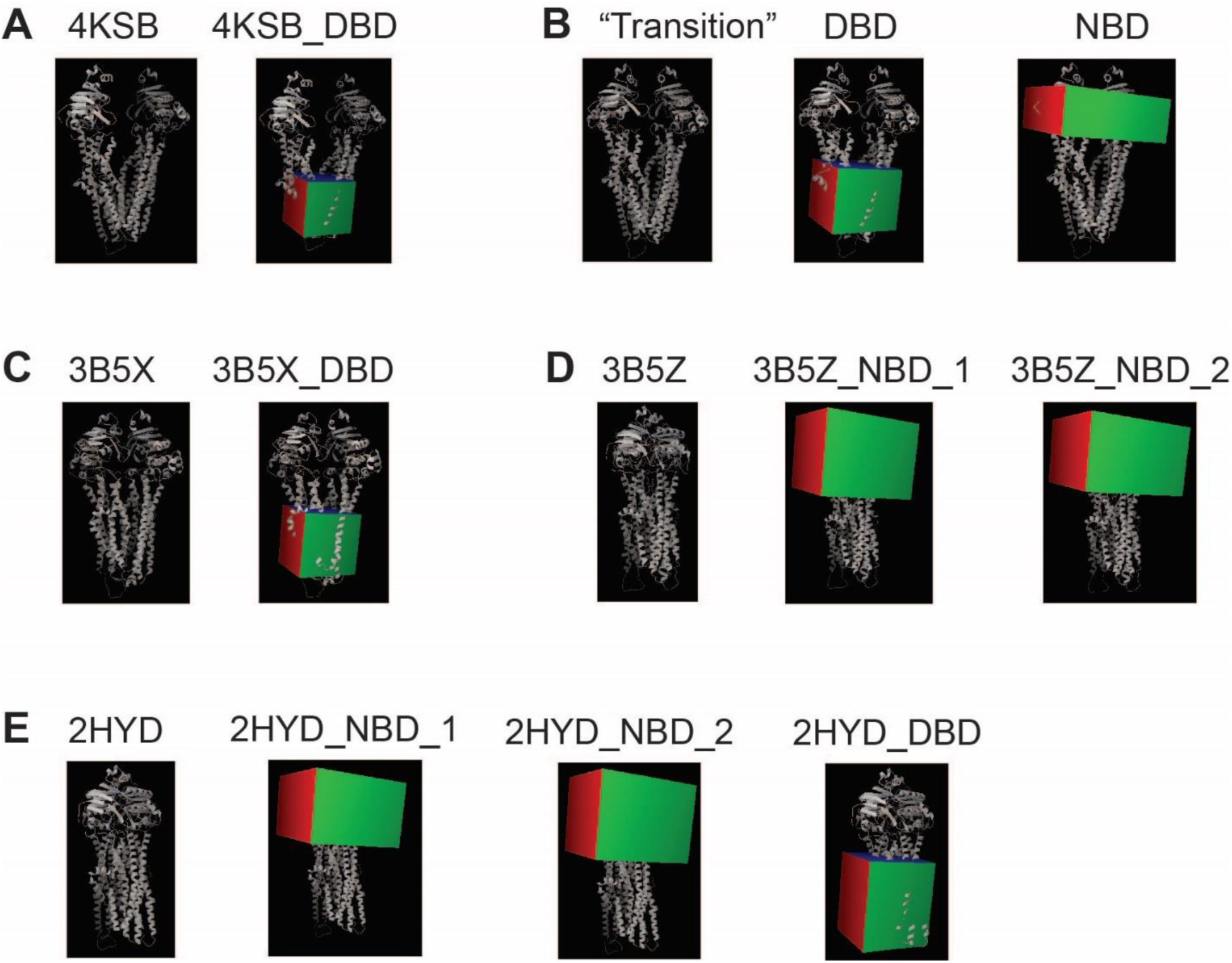
**Dock boxes and receptors for docking**. The conformations of the human P-gp model, and the corresponding dock boxes used, are shown in **A – E**. The first picture in each panel shows the receptor, with the PDB ID. The following pictures with red and green boxes show the docking boxes for the DBD or NBD of each receptor. Pictures generated using AutoDock. Note that the “Transition” structure is a conformation in-between 4KSB and 3B5X and is derived from TMD simulations performed for this study.

**S3 Fig.**
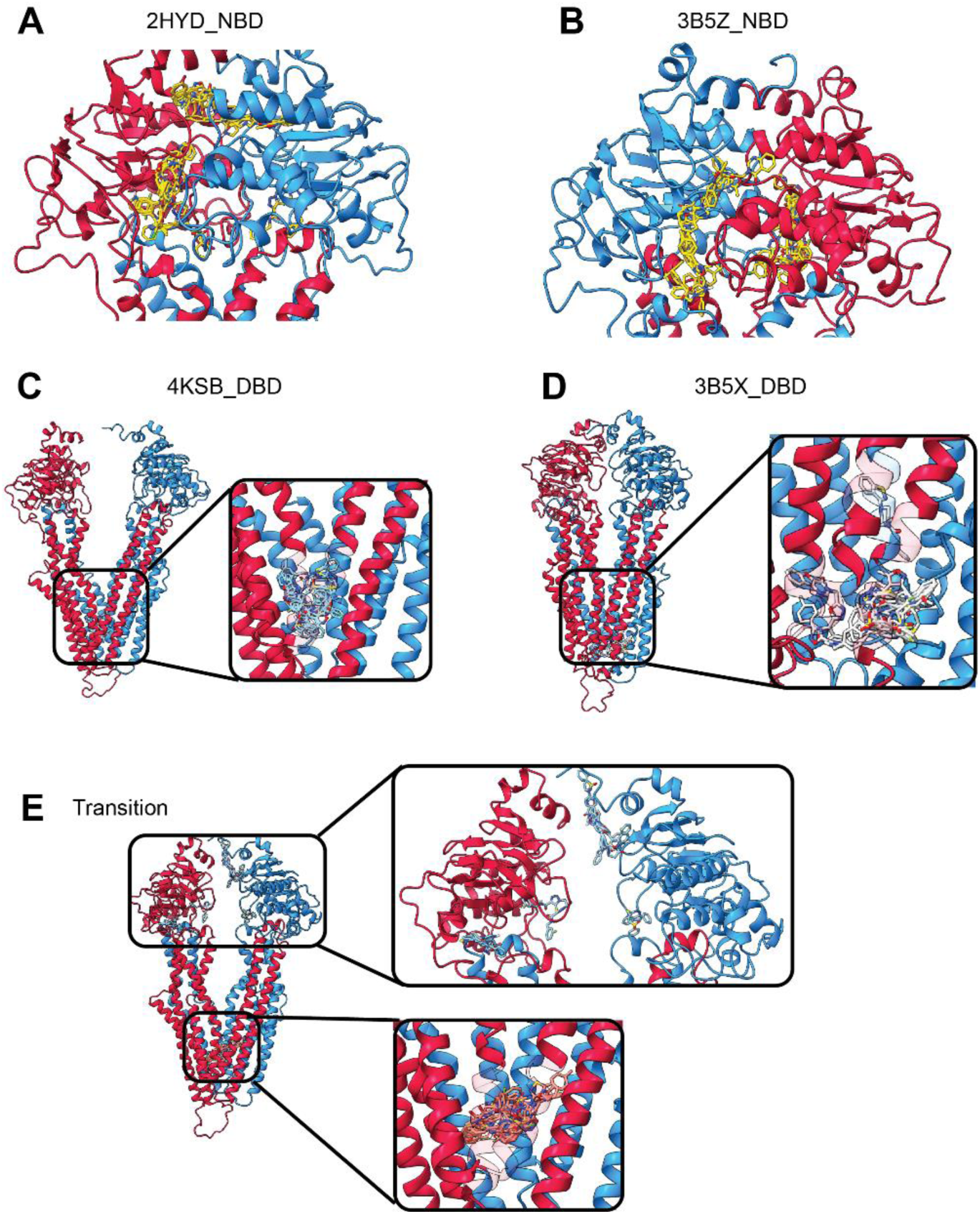
**Ligand docking locations to each P-gp structure**. The docking positions of the top hits to the NBDs of **A)** 2HYD and **B)** 3B5Z were mostly located within the ATP-binding site, with the exception of a few positions above the ATP-binding site in 2HYD. The docking positions of top hits to the DBDs of **C)** 4KSB and **D)** 3B5X were near the middle or the bottom of the drug binding region, respectively. Ligands were docked to both the **E, top box)** NBDs and **E, bottom box)** DBDs of the “Transition” structure, which was generated from MD simulation trajectories that transitioned the protein from the 4KSB to the 3B5X conformation. It is notable that several hits were predicted to bind near the ATP-binding sites, even though, when the NBDs are disengaged, the catalytic configuration for ATP hydrolysis is incomplete.

**S4 Fig.**
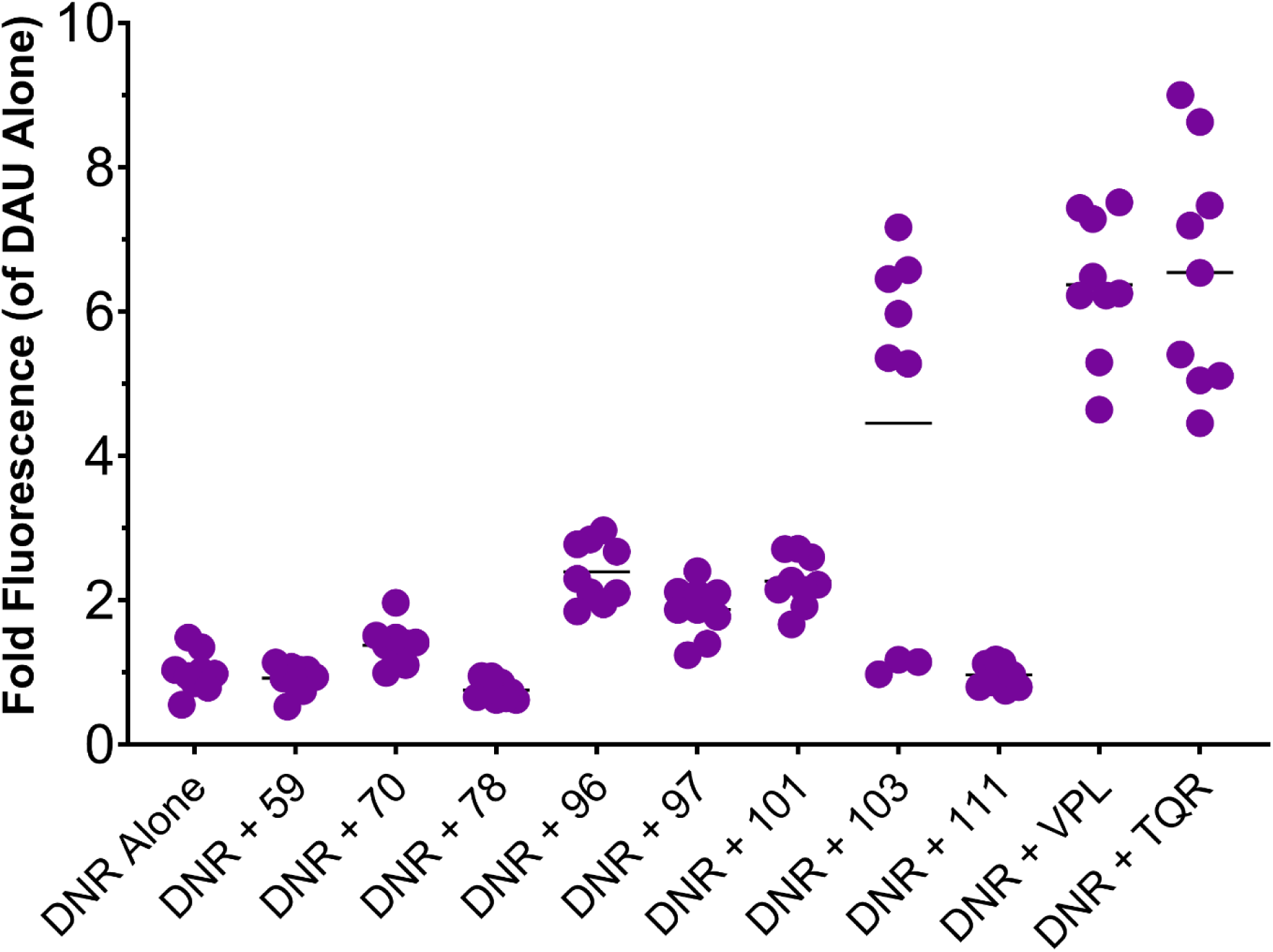
Alternative representation of daunorubicin fluorescence testing of novel inhibitors. An alternative representation of the data in Fig 5. Fold change in fluorescence of the P-gp substrate Daunorubicin (DAU) in the presence or absence of experimental compounds or known P-gp modulators. Cells were treated with 10 µM compound in the presence or absence of 10 µM DAU. After washing and lysing the cells, the intracellular DAU fluorescence was measured and expressed as a fold change relative to the fluorescence of DU145-TXR cells treated with DAU alone. The P-gp inhibitors, VPL and TQR, were included as positive controls for P-gp inhibition, and compound 59 was included as a negative control for P-gp inhibition. Three samples per trial, three independent trials. Significance was determined using a Student’s T test of the mean by comparing fluorescence of DAU and compound to that of DAU alone; P > 0.05 = N.S., P < 0.05 = *, P < 0.01 = **, P < 0.001 = ***, P < 0.0001 = ****. Note that for compound 78, the significance denotes a significant decrease in fluorescence.

**S1 Table.** Known P-gp Inhibitors and Predicted Chemical Properties. . A non-exhaustive list of known P-gp inhibitors and their selected chemical properties [64, 65]. Clinical trial outcomes were adapted from [10, 11]. Compounds were evaluated using the SWISS-ADME server [54]. RO5: Rule of 5, MW: Molecular Weight, TPSA: Topological Polar Surface area, LogP: Average of 5 calculations of LogP_oct/wat_, LogS: Average of 3 calculations to determine aqueous solubility – Insoluble, Poorly soluble (Poor), Moderately Soluble (Moderate), Soluble. “Predicted P-gp Substrate” - predicted by SWISS-ADME to be a transport substrate of P-gp. “Known P-gp Substrate” – In addition to being P-gp inhibitors, these molecules are: known transport substrates of P-gp, known non-transport substrates of P-gp, unknown whether it is a transport substrate of P-gp [66–68]. If the molecule is designated in a specific generation of P-gp inhibitors - First Generation P-gp inhibitors, Second generation P-gp inhibitors, Third generation P-gp inhibitors.

| Generation of P-gp Inhibitors |  | Compound | MW (g/mol) | Clinical Trial Result | # RO5 Violations | TPSA (Å<sup>2</sup>) | Log P<sub>oct/wat</sub> | LogS | Predicted P-gp Substrate | P-gp Substrate ? |
| --- | --- | --- | --- | --- | --- | --- | --- | --- | --- | --- |
|  | 1<sup>st</sup> | Verapamil | 455 | Fail | 0 | 64 | 4.45 | Moderate | Yes | Yes |
|  |  | Reserpine | 609 |  | 2 | 118 | 3.52 | Poor | Yes | No |
|  |  | Cyclosporin A | 1203 | Fail | 2 | 279 | 2.38 | Poor | Yes | Yes |
|  |  | Quinidine | 324 | Fail | 0 | 46 | 2.81 | Soluble | No | Yes |
|  |  | Amiodarone | 645 | Fail | 2 | 43 | 6.49 | Poor | Yes | Yes |
|  | 2<sup>nd</sup> | Valspodar | 1215 | Fail | 2 | 276 | 2.26 | Poor | Yes | No |
|  |  | Elacridar | 564 | Fail | 1 | 93 | 4.93 | Poor | No | Yes |
|  |  | Dexverapamil | 455 | Fail | 0 | 64 | 4.45 | Moderate | Yes | No |
|  | 3<sup>rd</sup> | Tariquidar | 647 | Fail | 1 | 111 | 5.2 | Poor | No | Yes |
|  |  | Zosuquidar | 528 | Fail | 1 | 49 | 4.64 | Poor | No | Yes |

**S2 Table.** Parameters of dock boxes used for docking screens.

| Receptor Name | PDB source | Search Area | Center coordinates |  |  | Size (Å) |  |  |
| --- | --- | --- | --- | --- | --- | --- | --- | --- |
|  |  |  | X | Y | Z | X | Y | Z |
| 2HYD_DBD_1 | 2HYD | DBD | -7.7 | 0.0 | 3.2 | 50 | 50 | 60 |
| 2HYD_NBD_1 | 2HYD | NBD | -4.3 | 2.5 | 70.1 | 76 | 64 | 58 |
| 2HYD_NBD_2 | 2HYD | NBD | -3.9 | 1.7 | 70.1 | 78 | 76 | 50 |
| 3B5Z_NBD_1 | 3B5Z | NBD | -4.3 | 2.5 | 70.1 | 76 | 64 | 58 |
| 3B5Z_NBD_2 | 3B5Z | NBD | -3.9 | 1.7 | 70.1 | 78 | 76 | 50 |
| 3B5X_DBD | 3B5X | DBD | -0.9 | 1.7 | 2.6 | 44 | 40 | 44 |
| Transition_NBD |  | NBD | -2.5 | 3.8 | 61.4 | 78 | 76 | 30 |
| Transition_DBD |  | DBD | -0.9 | 1.7 | 2.6 | 50 | 46 | 44 |
| 4KSB_DBD | 4KSB | DBD | -1.4 | 0.8 | -2.8 | 40 | 40 | 40 |

**S3 Table.** Ratio of estimated affinities (K_D_^estimate^) for the top hits. Ratios are calculated as the estimated DBD affinity (K_D_^DBD^) divided by the estimated NBD affinity (K_D_^NBD^). Ratios are listed by compound, and then organized by receptor structure. Three DBD docking boxes and five NBD boxes were used (**Error! Reference source not found.**). Each NBD of the 2HYD and 3B5Z structures was given its own docking box, hence the labels “nbd_1” and “nbd_2”. Molecules with a high ratio of DBD/NBD affinities are the most desirable, as this indicates that the K_D_^estimate^ to the DBDs is much larger (and thus lower affinity) than the K_D_^estimate^ to the NBDs.

| SMU ID | 4KSB DBD / NBD |  |  |  |  |
| --- | --- | --- | --- | --- | --- |
|  | 2hyd_nbd_1 | 2hyd_nbd_2 | 3b5z_nbd_1 | 3b5z_nbd_2 | transition_nbd |
| 70 | 3.3 | 57.3 | 3.9 | 3.3 | 6.4 |
| 78 | 1.0 | 1.0 | 1.2 | 0.2 | 0.4 |
| 96 | 0.3 | 0.4 | 0.0 | 0.0 | 0.0 |
| 97 | 2.8 | 0.7 | 1.0 | 0.1 | 0.1 |
| 101 | 2.0 | 40.9 | 5.4 | 2.8 | 2.8 |
| 103 | -- | 12.6 | 3.9 | 0.4 | 1.7 |
| 111 | 0.7 | 0.5 | 0.7 | 0.1 | 0.7 |
| 122 | 14.9 | 7.6 | 17.6 | 3.9 | 2.3 |
| 124 | 7.6 | 57.3 | 1.7 | 5.4 | 1.2 |
| SMU ID | 3B5X DBD / NBD |  |  |  |  |
|  | 2hyd_nbd_1 | 2hyd_nbd_2 | 3b5z_nbd_1 | 3b5z_nbd_2 | transition_nbd |
| 70 | 6.4 | 112.6 | 7.6 | 6.4 | 12.6 |
| 78 | 3.9 | 3.9 | 4.6 | 0.8 | 1.4 |
| 96 | 3.9 | 6.4 | 0.2 | 0.1 | 0.1 |
| 97 | 6.4 | 1.7 | 2.3 | 0.3 | 0.2 |
| 101 | 2.3 | 48.4 | 6.4 | 3.3 | 3.3 |
| 103 | -- | 14.9 | 4.6 | 0.5 | 2.0 |
| 111 | 6.4 | 4.6 | 6.4 | 1.2 | 6.4 |
| 122 | 7.6 | 3.9 | 9.0 | 2.0 | 1.2 |
| 124 | 9.0 | 67.9 | 2.0 | 6.4 | 1.4 |
| SMU ID | Transition DBD / NBD |  |  |  |  |
|  | 2hyd_nbd_1 | 2hyd_nbd_2 | 3b5z_nbd_1 | 3b5z_nbd_2 | transition_nbd |
| 70 | 0.8 | 14.9 | 1.0 | 0.8 | 1.7 |
| 78 | 1.7 | 1.7 | 2.0 | 0.4 | 0.6 |
| 96 | 4.6 | 7.6 | 0.3 | 0.2 | 0.1 |
| 97 | 2.3 | 0.6 | 0.8 | 0.1 | 0.1 |
| 101 | 1.2 | 24.7 | 3.3 | 1.7 | 1.7 |
| 103 | -- | 1.4 | 0.4 | 0.0 | 0.2 |
| 111 | 0.8 | 0.6 | 0.8 | 0.2 | 0.8 |
| 122 | 1.4 | 0.7 | 1.7 | 0.4 | 0.2 |
| 124 | 4.6 | 7.6 | 0.3 | 0.2 | 0.1 |

**S4 Table.**
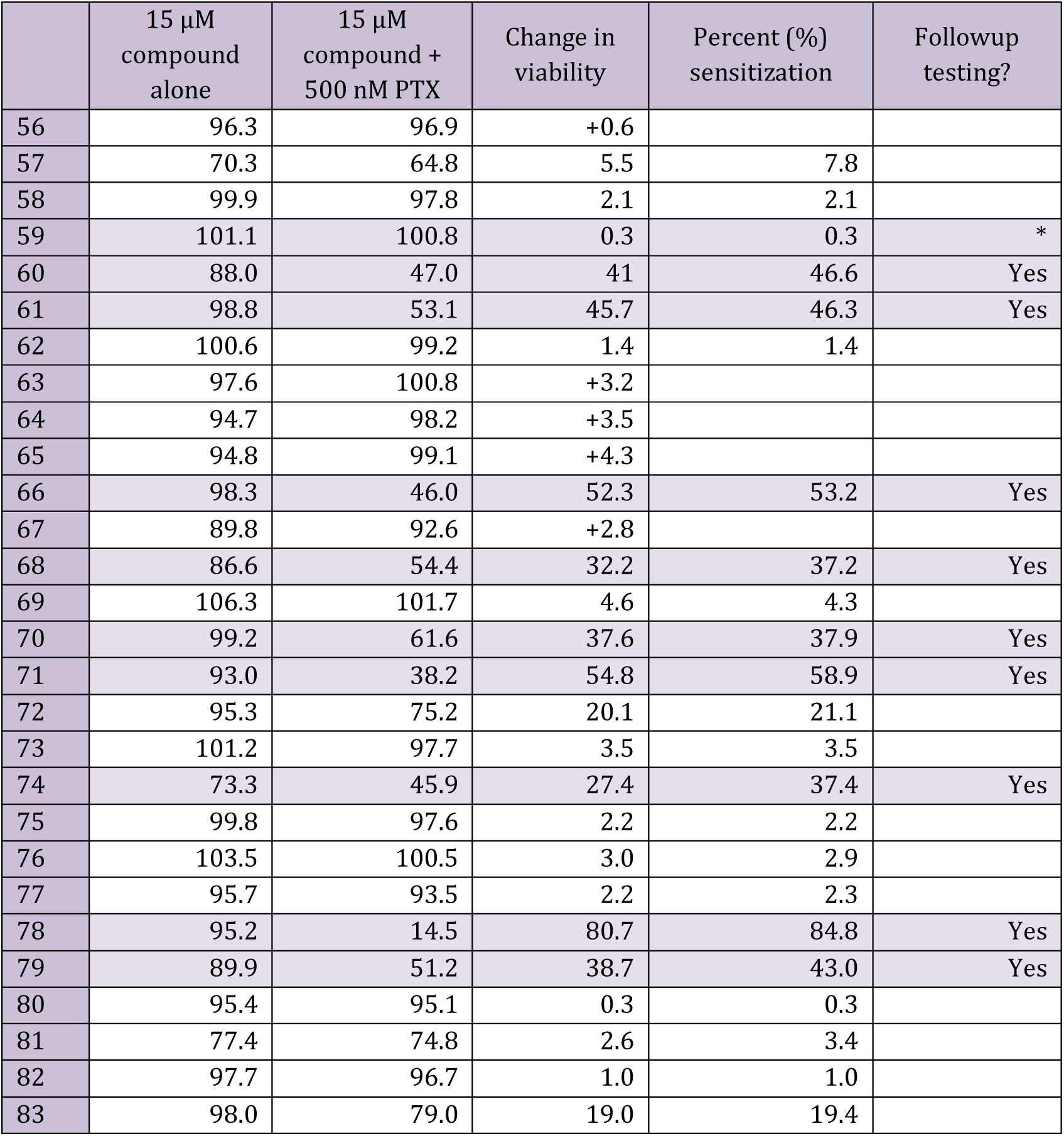
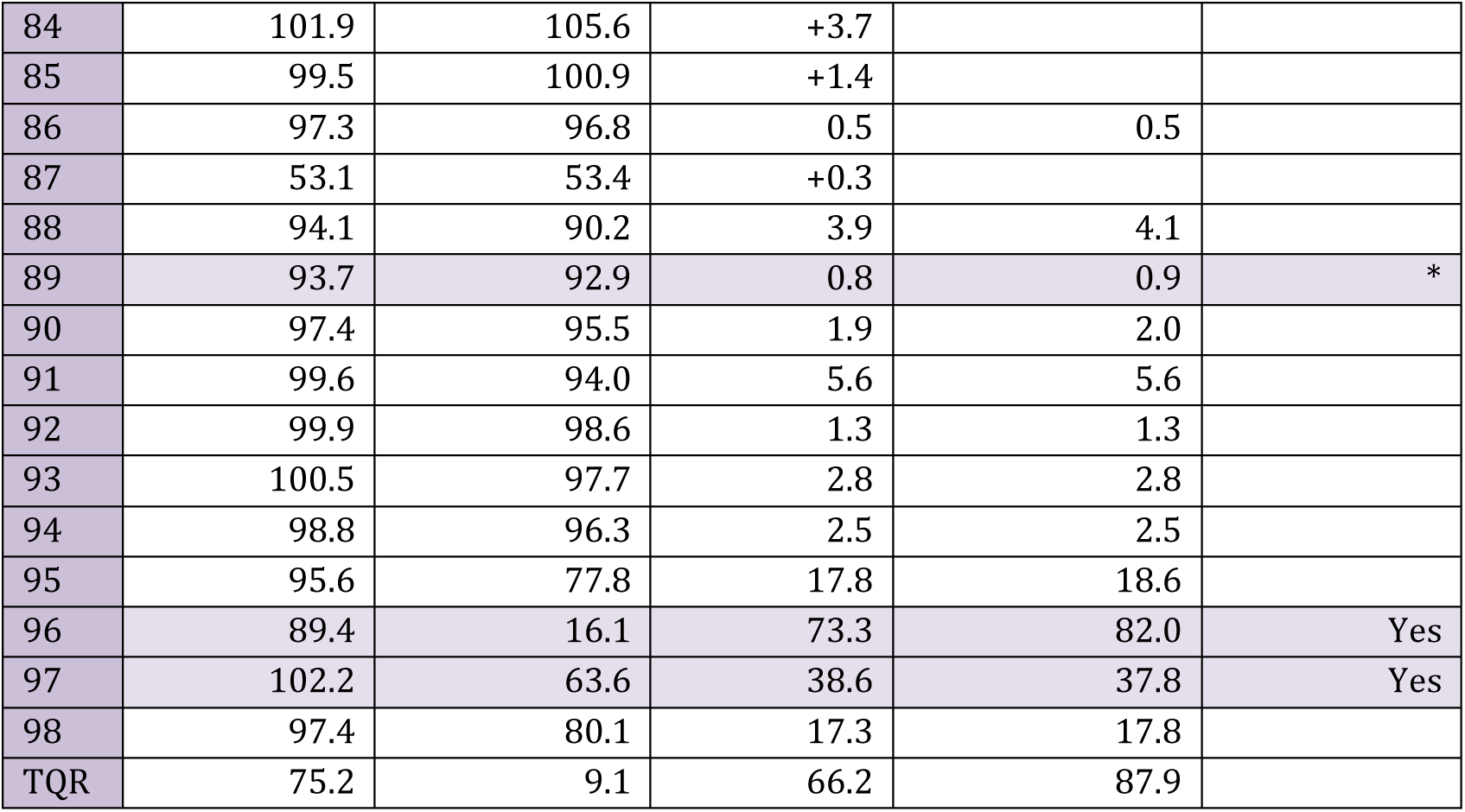
Pre-screening of compounds 56 – 98 against DU145-TXR cells with resazurin assays. . DU145-TXR cells were incubated with 15 µM compound with or without 500nM PTX for 48 hrs; survival was subsequently determined with the Resazurin viability assay [37]. Data represent the mean of two separate experiments performed in triplicate, and shows viability of cells treated with 15 µM compound alone, 15 µM compound and 500 nM PTX, and the difference between survivability measurements of each treatment. Percent re-sensitization is defined as the percent change in viability between cells treated with compound and PTX, versus cells treated with PTX alone. In some instances, the assay reported an increase in viability with compound + PTX, and no percent sensitization is reported. ‘Follow-up Testing’ compounds were re-assessed with MTT assays (Figure 1). Compounds 59 and 89 (*, ‘Follow-up Testing’) were used as negative controls in follow-up MTT assays to test for consistency of results, e.g. to confirm that molecules eliminated in Resazurin assays were justifiably eliminated from later screening with MTT assays. Tariquidar was included as a positive control for P-gp inhibition.

**S5 Table.**
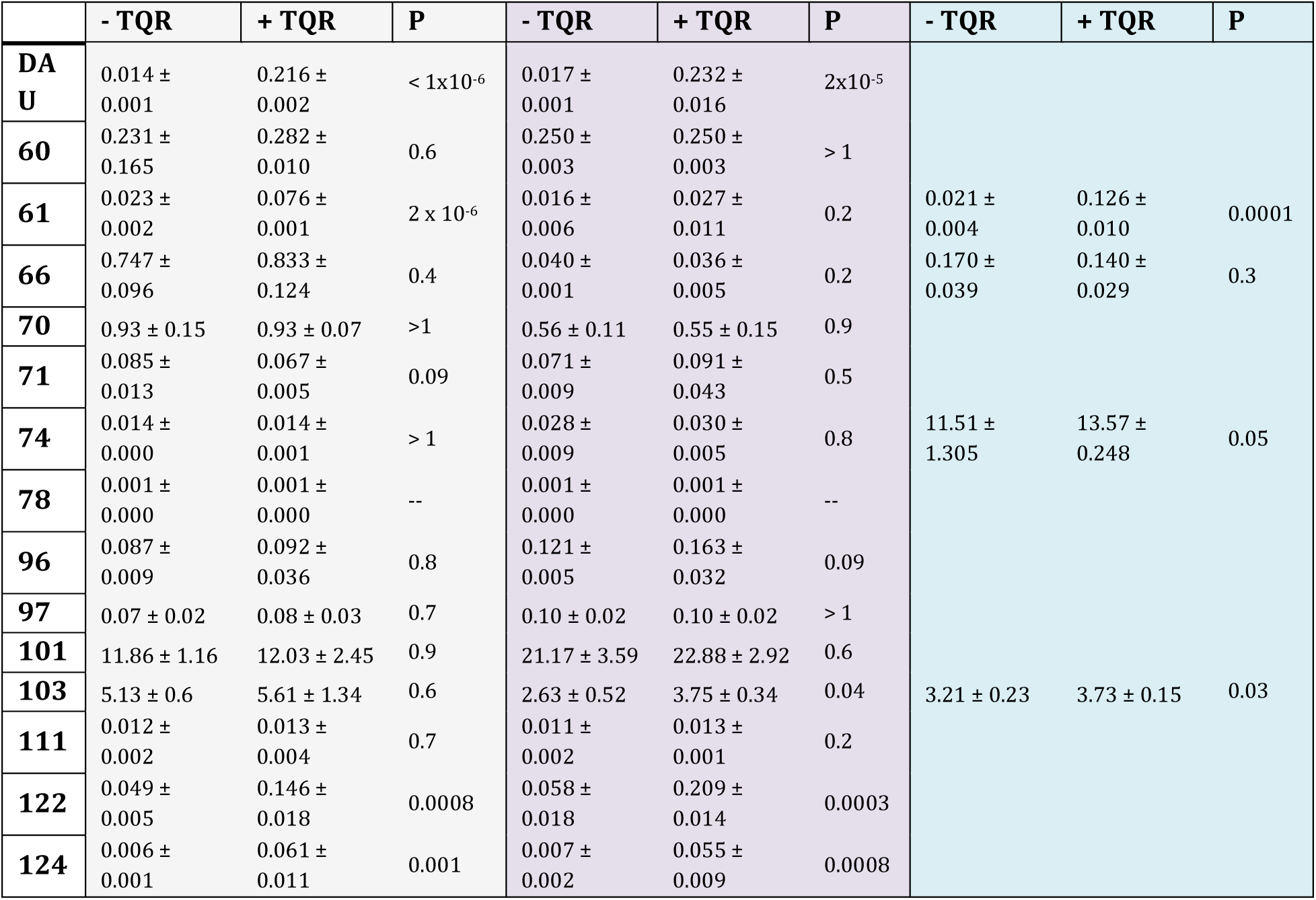
Normalized ratio of compound to internal standard using LC-MS/MS. . DU145-TXR cells were exposed to 5 µM compound with or without 500 nM tariquidar (TQR) as described in [8]. Samples were prepared in triplicate and two or three independent trials were performed; data represent the mean ± one standard deviation (std. dev). In contrast to the methods used in [8], these LC-MS/MS trials measured the normalized ratio of analyte (i.e. compound) to the internal standard (i.e. unique performance of mass spectrometer on date of analysis); thus the data represent the relative quantification of compound per sample and explain the variability between independent trials (see Methods).

**S6 Table.**
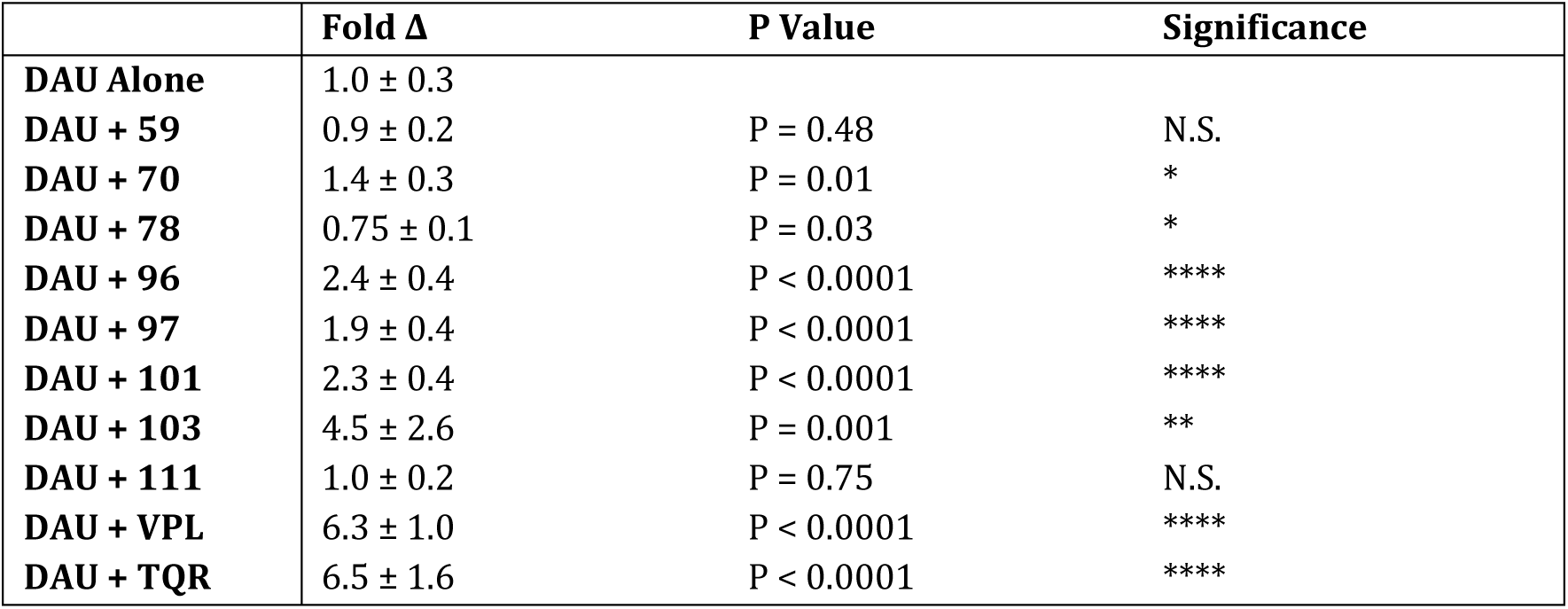
Fold Accumulation of Daunorubicin fluorescence in DU145-TXR cells. Here we show the fold fluorescence of the P-gp substrate Daunorubicin (DAU) in the presence or absence of experimental compounds or known P-gp modulators. Cells were treated with 10 µM compound in the presence or absence of 10 µM DAU, after which cells were washed and lysed; the resultant DAU fluorescence was measured on the Cytation 5 (excitation/emission 488 nm / 575 nm). The resultant DAU fluorescence is expressed as a fold change relative to the fluorescence of DU145-TXR cells treated with DAU alone. The P-gp inhibitors VPL and TQR were included as positive controls for P-gp inhibition, and compound 59 was included as a negative control for P-gp inhibition. Three samples per trial, three independent trials. Statistical significance determined using GraphPad Prism, Student’s T test of the mean, by comparing the mean fluorescence of DAU + compound to that of DAU alone. Significance was determined using a Student’s T test of the mean; P > 0.05 = N.S., P < 0.05 = *, P < 0.01 = **, P < 0.001 = ***, P < 0.0001 = ****.

**S7 Table.**
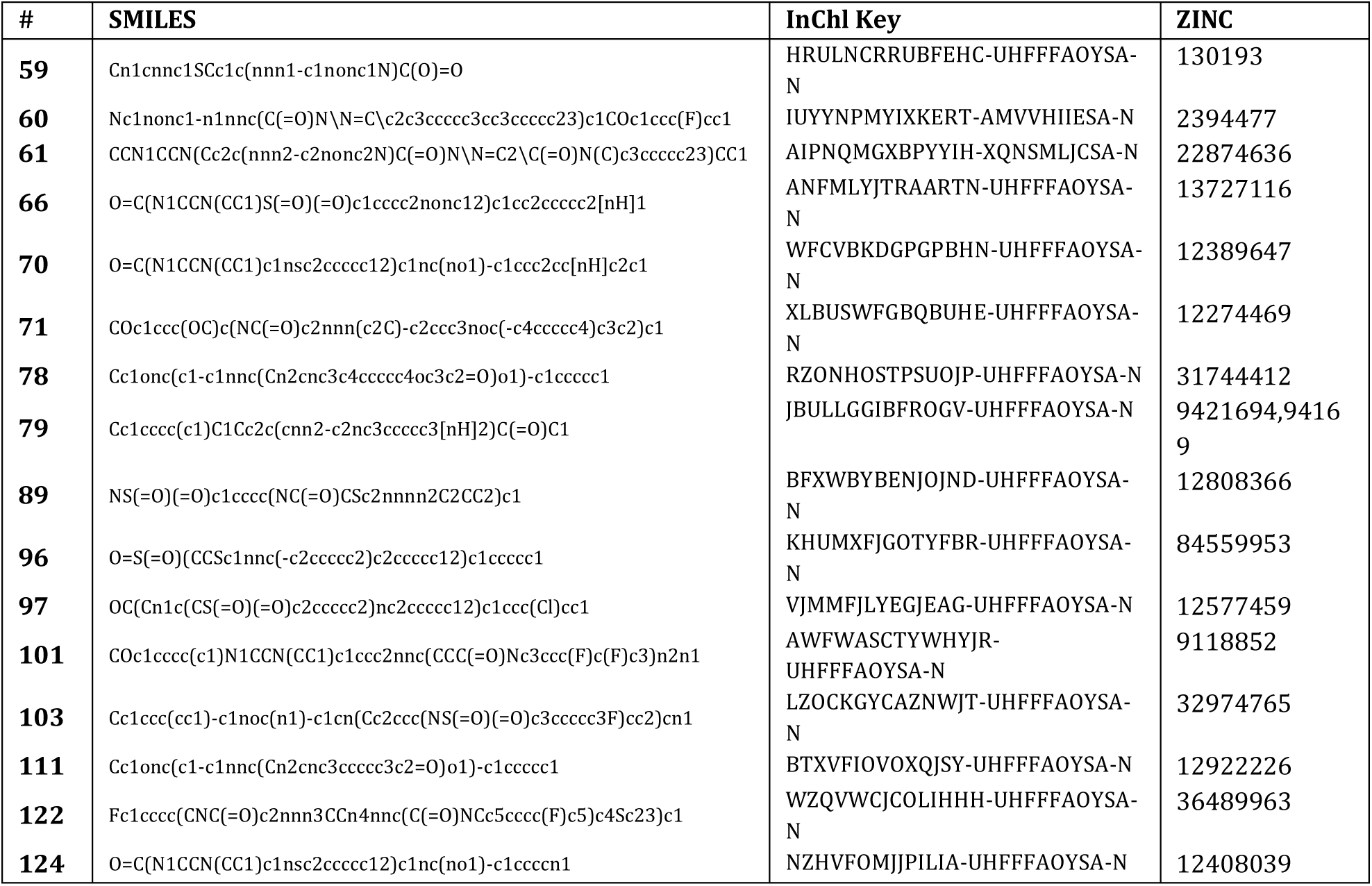
Alternative identifiers for molecules identified in this study. ZINC IDs are shown if available. Molecular structures are translated into SMILES, and the corresponding InChl keys are provided as well.

